# Direct RNA-binding by MYCN mediates feedback from RNA processing to transcription control

**DOI:** 10.1101/2023.08.16.553474

**Authors:** Dimitrios Papadopoulos, Stefanie Anh Ha, Daniel Fleischhauer, Leonie Uhl, Katharina Schneider, Ivan Mikicic, Timothy J. Russell, Annika Brem, Omkar Rajendra Valanju, Peter Gallant, Christina Schuelein-Voelk, Hans Michael Maric, Petra Beli, Gabriele Büchel, Seychelle M. Vos, Martin Eilers

## Abstract

The MYCN oncoprotein broadly binds active promoters in a heterodimer with its partner protein MAX. MYCN also interacts with the nuclear exosome, a 3’-5’ exoribonuclease complex, suggesting a function in RNA metabolism. Here we show that MYCN forms stable high molecular weight complexes with the exosome and multiple RNA-binding proteins. In cells, MYCN binds to thousands of intronic RNAs; recombinant MYCN directly binds RNA via a short, highly conserved sequence termed MYCBoxI. Perturbing exosome function results in global re-localization of MYCN from promoters to intronic RNAs. At promoters, MYCN is then replaced by the MNT(MXD6) repressor protein, which inhibits MYCN-dependent transcription. MYCN promotes the degradation of its bound introns via the nuclear exosome targeting (NEXT) complex. Our data demonstrate that MYCN is an RNA-binding protein that regulates nascent transcript turnover and show that competition between its RNA– and DNA-bound states links the dynamics of the MYCN/MAX/MXD network to mRNA processing.

## Introduction

The members of the *MYC* oncogene family, *MYC*, *MYCN* and *MYCL*, drive oncogenesis in a wide variety of human tumors and expression of each paralog is associated with a characteristic spectrum of tumor entities^1,2^. Expression of MYCN is found, for example, in subentities of adult prostate and small cell lung carcinoma as well as in childhood medulloblastoma and neuroblastoma. A paradigm example are *MYCN*-amplifications in neuroblastoma that drive the development of a distinct subentity with a particularly poor prognosis^3^.

MYCN, like its paralogue MYC, is a DNA-binding transcription factor that can affect the expression of a broad ensemble of target genes^4–6^. To bind to DNA, MYC proteins form a heterodimer with MAX via a conserved helix-loop-helix leucine zipper domain. Through this domain, MAX can also form homodimers and heterodimers with MXD proteins, which compete with MYCN for binding to MAX^7,8^. The different heterodimers have antagonistic functions: whereas MYCN/MAX heterodimers activate transcription, MAX/MAX homodimers are considered neutral and MXD/MAX repressors of transcription^8^. To activate transcription, MYC and MYCN interact with TRRAP, a scaffold protein of several histone acetylase complexes^9–12^. Conversely, MXD proteins interact with SIN3A, an adaptor protein of histone deacetylase complexes^13,14^, hence the primary functional output of the MYCN/MAX/MXD network are widespread changes in histone acetylation at promoters and enhancers^2,15–17^. Remarkably, several MXD proteins are co-expressed with MYCN or MYC in highly proliferating cells, and the mechanisms that control the dynamics of the network in proliferating cells are unknown. Proteomic analyses show that the MYC and MYCN interactomes are surprisingly complex and that MYCN is not only involved in protein complexes that function early in transcription at promoters, but also in transcription termination, DNA repair and protein turnover^18–26^. Functional screening in neuroblastoma cells shows that the proliferation of *MYCN*-amplified neuroblastomas depends on MYCN interaction partners that differ from those of MYC-driven neuroblastoma^24^. Specifically, MYCN-driven tumor cells show an enhanced dependence on the nuclear exosome, a 3’-5’ RNA exonuclease that degrades multiple non-coding RNA species shortly after their synthesis^24,27^ and can facilitate both restart and termination of stalled RNA polymerase II (RNAPII)^28,29^. Via association with the exosome, MYCN prevents conflicts of the replication fork with stalled RNAPII and maintains genomic stability during S phase^24^. The observation that a MYCN-exosome complex has a specific function during S phase supports previous findings showing that MYCN forms distinct complexes during cell cycle progression^22^. Since the nuclear exosome is not associated with DNA or promoters, the data also raised the possibility that MYCN participates in RNA-binding complexes independent of its association with DNA and may have a role in RNA metabolism. Here we show that MYCN binds directly to RNA mainly via a conserved sequence element in the N-terminal region of MYCN termed MYCBoxI. In cells, MYCN is bound to hundreds of intronic RNAs and RNA-binding of MYCN controls the dynamics and the transcriptional output of the MYCN/MAX/MXD network. When bound to RNA, MYCN is part of the <u>n</u>uclear <u>ex</u>osome targeting (NEXT) complex which directs the exosome to non-polyadenylated transcripts thereby promoting their degradation.

## Results

### MYCN forms stable complexes with the nuclear exosome and other RNA-binding proteins

We initially explored the possibility that MYCN forms complexes in S phase that are distinct from those present in G1 and G2 phases. To do so, we loaded *MYCN*-amplified IMR-5 neuroblastoma cell lysates that were prepared from either asynchronously growing cells or from cells synchronized in S phase with double-thymidine block on pre-calibrated glycerol gradients (Figures S1A,B). Inspection of the sedimentation patters showed that in asynchronously growing cells the majority of MYCN fractionated at a molecular weight compatible with it being a monomer or forming a heterodimeric complex with MAX (Figures S1C,D). In contrast, a large fraction of MYCN, but not MAX, recovered from cells synchronized in S phase formed much larger complexes with range of molecular weights of one megadalton and higher (Figures S1C,D). Multiple subunits of the nuclear exosome as well as other RNA-binding proteins (RBPs) were recovered in MYCN immunoprecipitates from high molecular weight complexes of S phase cells, but to a much lesser degree from asynchronous cells (Figure S1E). Collectively, these data show that MYCN forms stable high molecular weight complexes during S phase that contain the nuclear exosome.

To investigate the role of the nuclear exosome and its RNA substrates in MYCN complex formation, we used *MYCN*-amplified IMR-32 neuroblastoma cells that stably express a doxycycline-inducible shRNA targeting EXOSC10, one of the two catalytic subunits of the exosome (Figure 1A)^27^. Nascent RNA sequencing (4sU-seq) showed that depletion of EXOSC10 led to the accumulation of multiple short-lived RNA species, for example promoter upstream transcripts (“PROMPT”), consistent with its role in RNA degradation (Figures 1B,C and Figure S2A). Gradient centrifugation showed that depletion of EXOSC10 caused the accumulation of MYCN in high molecular weight complexes (Figure 1D). As in S phase-synchronized cells, MAX migrated at a low molecular weight corresponding to a heterodimer with MYCN or MXD-family partner proteins in both control and EXOSC10-depleted cells, suggesting that a fraction of MYCN accumulates without MAX in high molecular weight complexes (Figures 1D,E). Proteomic analysis showed a fundamental change in the MYCN interactome upon EXOSC10 depletion (Figure 1F). In particular, we observed a strong decrease in interactions with TRRAP and NuA4, the major MYCN-associated histone acetylase complex for which TRRAP serves as scaffold^22^, as well as an increase in association of MYCN with multiple RNA-binding proteins and the PAF1 complex (Figures 1F,G). Proximity ligation assays measuring the proximity of MYCN to other proteins confirmed these results by documenting a decrease in association of MYCN with MAX, with TRRAP and the catalytic subunit of the NuA4 complex, KAT5 (TIP60), as well as with TOP2B, a transcriptional coactivator of MYCN^30^, upon EXOSC10 depletion (Figures S2B,C).

**Figure 1:**
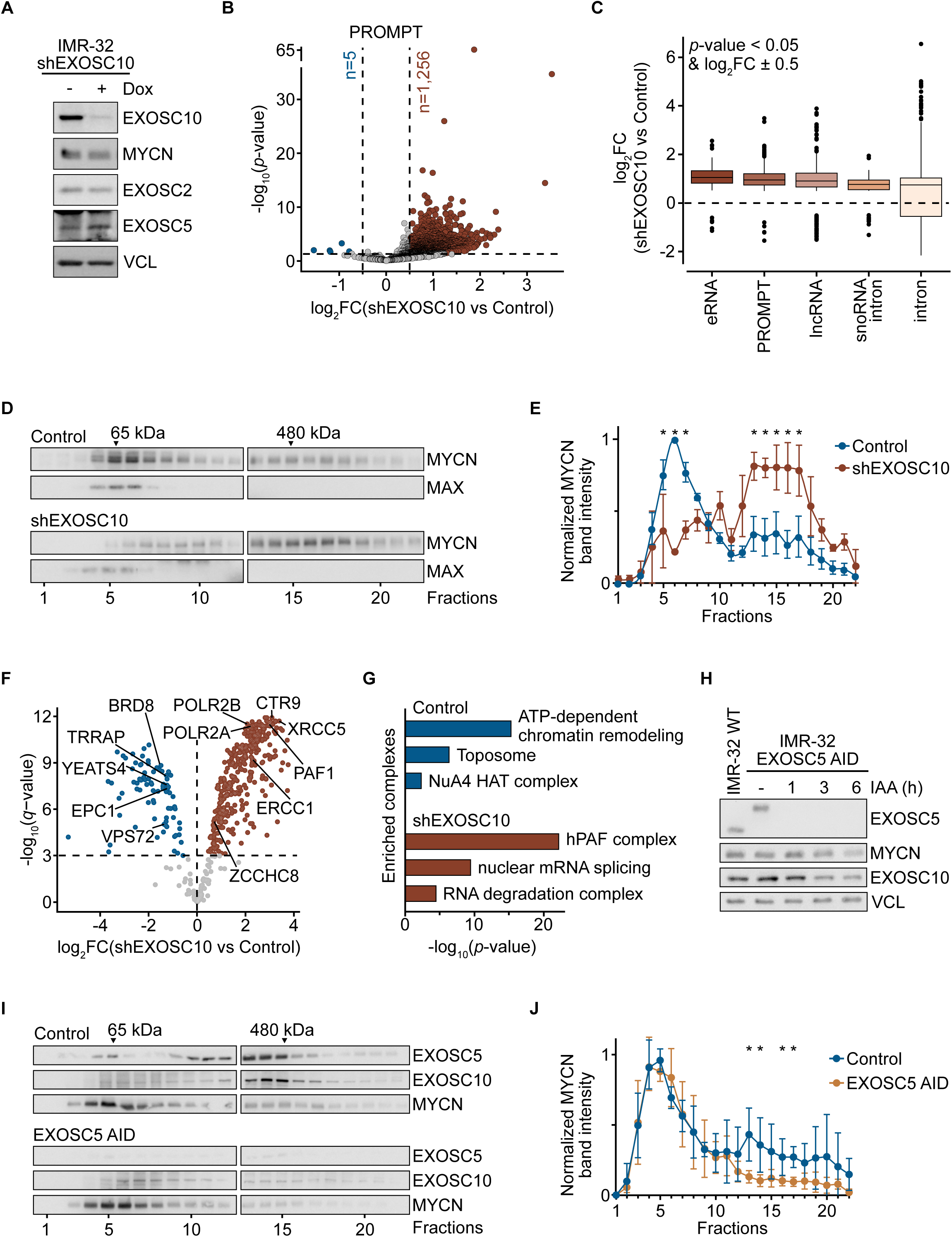
MYCN forms high molecular weight complexes with the nuclear exosome and other RNA-binding proteins. (**A**) Immunoblot of IMR-32 cells stably expressing an shRNA against EXOSC10 treated for 72 h with doxycycline (Dox) (1 µg/ml) or EtOH as control. Vinculin (VCL) was used as a loading control (n=3; n indicates independent biological replicates). **(B)** Volcano plot showing log_2_ fold-change (FC) (x-axis) and statistical significance (y-axis) of 4sU-labeled RNA levels between shEXOSC10 and control IMR-32 cells, treated as in (A). Each point represents an annotated promoter upstream transcript (PROMPT). PROMPTs with *p*-value < 0.05 and log_2_FC > 0.5 or < –0.5 were considered significantly increased (brown) or decreased (blue), respectively (n=3). **(C)** Log_2_FC comparing 4sU-labeled RNA levels of the indicated groups of non-coding RNAs between shEXOSC10 and control IMR-32 cells from the same experiment as in (B). Only RNAs passing shown thresholds were included in the plots. Lines within box plots indicate median, outliers shown as dots. **(D)** Immunoblot of endogenous MYCN and MAX sedimentation patterns in 10-30% glycerol gradients from IMR-32 cells treated as in (A) (n=3). **(E)** Quantification of the normalized MYCN band intensities from the immunoblots in (D). Dots indicate mean and error bars depict range of values. *P*-values were calculated using multiple unpaired t-tests (* indicates *p*-values < 0.05). **(F)** Volcano plot of MYCN protein interactions in IMR-32 cells following EXOSC10 depletion. Cells were treated as in (A), nuclear lysates were loaded on 10-30% glycerol gradients and MYCN immunoprecipitation (IP) was performed using pooled fractions 13-20 for both conditions. The x-axis displays the log_2_FC between shEXOSC10 and control cells (positive values indicate increased interactions with MYCN upon EXOSC10 loss). The y-axis shows statistical significance (*q*-value); a threshold of *q* < 0.001 was set for significant interactions (n=3). **(G)** Protein complexes exhibiting significantly altered associations with MYCN from interactome experiment shown in (F). **(H)** Immunoblot of IMR-32 wild-type (WT) and IMR-32 EXOSC5 AID (<u>a</u>uxin inducible <u>d</u>egron) cells treated with auxin Indole-3-acetic acid (IAA, 100 µM) for the indicated times. VCL was used as a loading control (n=3). **(I)** Immunoblot of MYCN, EXOSC10 and EXOSC5 sedimentation patterns in 10–30% glycerol gradients from nuclear lysates of IMR-32 EXOSC5 AID cells. In the control condition cells were treated with DMSO, in the EXOSC5 AID condition with IAA (100 µM) for 1.5 h (n=4). **(J)** Quantification of the normalized MYCN band intensities from the immunoblots shown in (I). Dots indicate mean and error bars depict range of values. *P*-values were calculated using multiple unpaired t-tests (* indicates p-values < 0.05).

To test whether the formation of high molecular weight complexes of MYCN depends on an intact exosome, we generated *MYCN*-amplified IMR-32 EXOSC5 AID cells, in which both alleles of the endogenous *EXOSC5* gene, which encodes a core structural subunit of the exosome^22^, were fused to a degron that is recognized by the plant-derived TIR1 ligase. In addition, we stably expressed the auxin-dependent TIR1 ligase in these cells. Addition of the auxin indole-3-acetic acid (IAA) to these cells led to a rapid degradation of EXOSC5, followed by a more gradual decrease in EXOSC10 levels (Figure 1H). The decrease in EXOSC10 levels is likely due to disintegration of the exosome as witnessed by a shift of EXOSC10 towards lower molecular weights in glycerol gradients (Figure 1I). Immunoblotting showed that MYCN protein levels are unaffected up to three hours after EXOSC5 degradation (Figure 1H). In contrast to EXOSC10 depleted cells, MYCN did not accumulate in high molecular weight complexes in EXOSC5 depleted cells as judged by glycerol gradient centrifugation (Figure 1I). Instead, we observed a significant decrease in the amount of MYCN present in such complexes, arguing that MYCN forms a significant portion of high molecular weight complexes with the nuclear exosome (Figures 1 I,J). We also observed that treatment of cell lysates with benzonase, which degrades both RNA and DNA, before centrifugation caused a minor decrease in the apparent molecular weights, but did not abolish formation of high molecular weight complexes of MYCN (Figure S2D). This is consistent with previous immunoprecipitation data of MYCN with exosome^24^, arguing that RNA contained in these complexes is either largely protected from benzonase digestion or not required for maintaining these complexes after they have formed. In summary, our findings indicate that cell cycle phase and accumulation of exosome targeted transcripts steer MYCN’s interactome away from DNA-centric complexes towards RNA-binding complexes.

### MYCN binds to intronic RNAs independently of DNA binding

These observations raised the possibility that MYCN, like the exosome, interacts with unstable or nascent RNAs in neuroblastoma cells. To test this hypothesis, we performed <u>e</u>nhanced <u>c</u>rosslinking and immuno<u>p</u>recipitation (eCLIP) experiments, in which proteins are crosslinked to RNA by UV irradiation and bound RNA molecules are subsequently recovered and sequenced^31^. These experiments documented a robust eCLIP signal for MYCN on multiple protein-coding genes (Figure 2A and Figure S3A). Notably, the genes where MYCN bound to RNA were distinct from the genes where MYCN typically binds DNA at promoters (Figure S3B).

**Figure 2:**
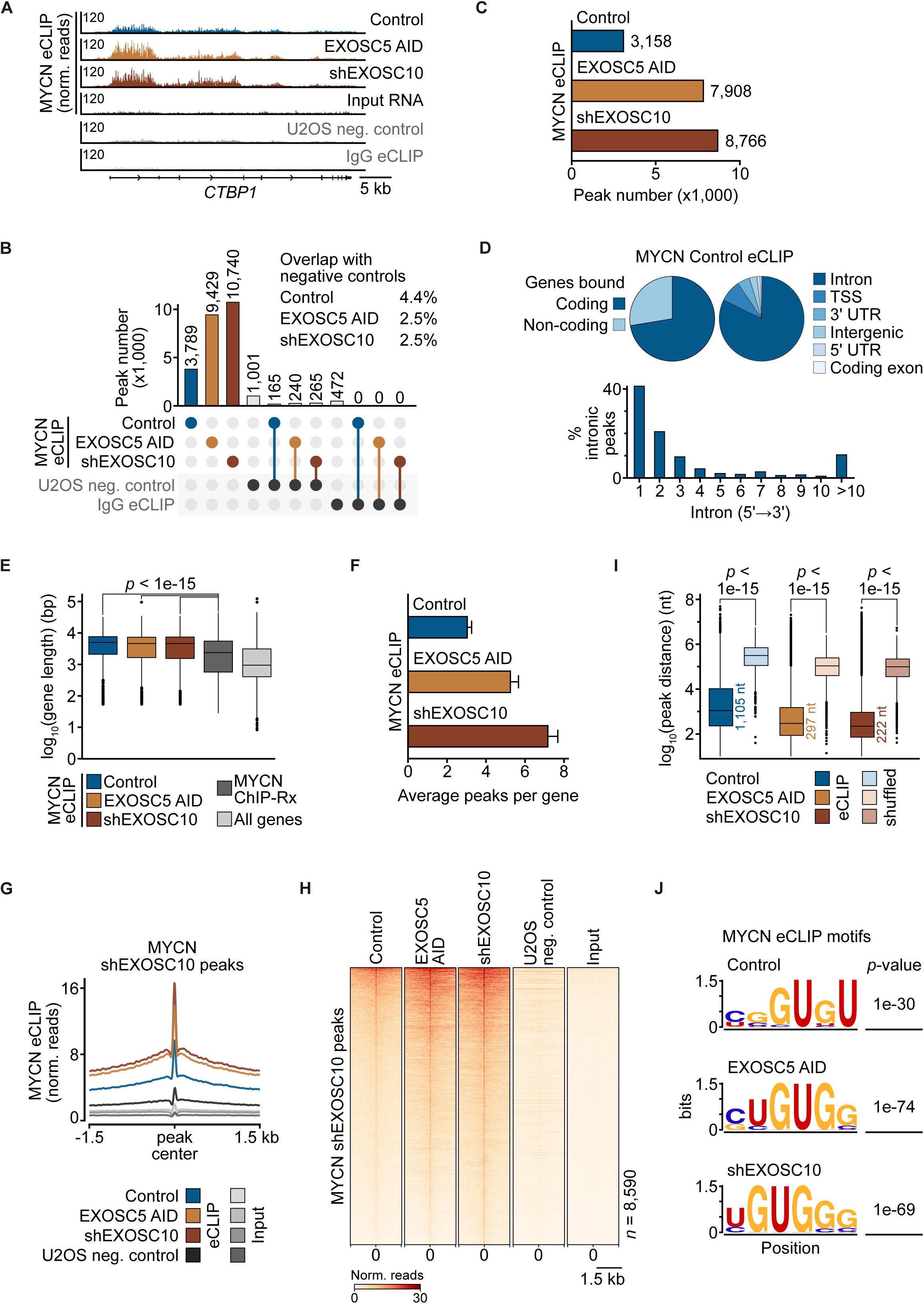
MYCN binds to intronic RNAs independently of DNA binding. (**A**) Read distribution of MYCN eCLIPs at the *CTBP1* gene. IMR-32 cells were treated with 72 h Dox (1 µg/ml) or 1.5 h IAA (100 µM) to induce EXOSC10 or EXOSC5 knock-down via shRNA or an AID, respectively. Control condition treated with EtOH. Input RNA corresponds to the control condition size-matched eCLIP input, U2OS neg. control refers to a MYCN eCLIP performed in U2OS cells that do not express MYCN, and IgG eCLIP was performed in untreated IMR-32 cells (n=2, all eCLIPs). **(B)** Overlap plot of high-confidence MYCN eCLIP peaks with U2OS neg. control peaks and IgG peaks. High-confidence peaks were defined by passing thresholds of log_2_FC > 3 over corresponding input and *p*-value < 0.001. (**C**) Number of high-confidence MYCN eCLIP peaks after removing all peaks overlapping with U2OS negative control and IgG eCLIP peaks and found within 1 kb of MYCN ChIP-Rx peak summits. MYCN ChIP-Rx was performed in untreated IMR-32 cells (n=3) and the replicate with the highest peak number was used for eCLIP filtering. (**D**) Peak distribution at coding and non-coding genes (top left) and at depicted gene regions (top right) for MYCN eCLIP control condition. Distribution of intronic MYCN eCLIP peaks from nearest to farthest from the transcription start site (bottom). (**E**) Lengths of all genes and of genes containing MYCN eCLIP or MYCN ChIP-Rx peaks. Lines within box plots indicate median, outliers shown as dots. *P*-values were calculated using Wilcoxon rank-sum test. (**F**) Average number of MYCN eCLIP peaks per gene in indicated conditions. Error bars show standard error of mean (n=2). (**G**) Trimmed (2%) average density blots of MYCN eCLIP and corresponding size-matched input reads in depicted conditions, centered around the called high-confidence MYCN shEXOSC10 peaks (n=8,590). Only reads in the sense strand are shown. Lines correspond to mean of biological duplicates for all eCLIPs, shaded regions depict standard error. (**H**) Heatmaps of trimmed (2%) reads of MYCN eCLIPs centered around MYCN shEXOSC10 peaks. Depicted input corresponds to the MYCN eCLIP control condition. All heatmaps were sorted based on descending mean reads in the MYCN eCLIP control condition. Only reads in the sense strand are shown. Shown are the means of biological duplicates for all eCLIPs. (**I**) Distances between the nearest MYCN eCLIP peak pairs in the depicted conditions. Shuffled datasets contain the same number of randomized intronic peaks as the corresponding eCLIP. Lines within box plots indicate median, outliers shown as dots. *P*-values were calculated using Wilcoxon rank-sum test. (**J**) *De novo* motif enrichment analysis based on the high-confidence intronic peaks from depicted MYCN eCLIPs against background of IMR-32 expressed intron sequences.

To test whether the interaction of MYCN with RNA depends on its interaction with the nuclear exosome, we repeated the eCLIP experiments upon depletion of EXOSC10 or EXOSC5 via shRNA or auxin induced degradation, respectively. In both conditions, the number of MYCN binding sites on RNA strongly increased. By performing stringent peak calling (> 8-fold enrichment over input RNA, *p*-value < 0.001) we identified 3,789 high-confidence binding sites for MYCN on RNA in untreated control cells. This number increased to 9,473 binding sites in EXOSC5 AID and 10,740 sites in EXOSC10 depleted cells (Figure 2B). No significant enrichment of specific RNA sequences relative to input RNA was observed in eCLIP experiments with control IgG or using the MYCN antibody in U2OS cells that do not express MYCN (U2OS neg. control), documenting the specificity of the signal (Figures 2A,B and Figures S3A,B). Furthermore, the binding sites defined by eCLIP showed low overlap (<10%) with MYCN ChIP-Rx peaks from the same cell line (Figure S3C). For all further analyses, we removed all eCLIP peaks overlapping with the peaks found in the negative control experiments or found within 1 kb of the MYCN ChIP-Rx peak summits and merged any bookended peaks, retaining more than 80% of all original peaks across all conditions (Figure 2C).

Annotation of the peaks to genomic regions showed that the majority of MYCN binding sites was localized in introns of protein-coding genes with a preference for 5’ proximal introns (Figure 2D). Loss of EXOSC5 or EXOSC10 did not significantly alter this distribution (Figure S3D). MYCN RNA-bound genes did not encode a functionally defined class of proteins as tested by gene set enrichment analysis (GSEA). However, they differed in length as MYCN RNA-bound genes were significantly longer than either all expressed genes or their DNA-bound counterparts hinting at a function for MYCN in the early synthesis of longer transcripts (Figure 2E). On individual genes, MYCN binding extended up to several kilobases and peak calling algorithms often identified multiple peaks per gene (Figure 2F). Density plots and heatmaps of eCLIP reads centered around the called peaks for MYCN showed that MYCN sharply binds RNA with a width of about 60 nucleotides (nt) and gradually decreases beyond this (Figures 2G,H). The distance between individual MYCN-bound peaks called on a single gene was not randomly distributed, but had a median distance of 1105 nt in untreated control, 297 nt in EXOSC5 and 222 nt in in EXOSC10 depleted cells, suggesting that multiple MYCN molecules bind to the same mRNA (Figure 2I). This is consistent with recent observations that a fraction of both MYCN and MYC exists in multimeric forms in cells^25,32^. *De novo* motif enrichment analyses performed against a background of expressed intronic sequences consistently showed that MYCN-bound RNA sequences were significantly enriched for UG-sequence motifs (Figure 2J and Figure S3E). G and U are the most commonly bound bases by RBPs^33^, suggesting that MYCN could potentially bind RNA in complex with additional RBPs, such as the NEXT complex of the RNA exosome. This will be discussed further below. Collectively, the data show that MYCN is bound to the intronic RNA of protein-coding genes.

### Direct binding of MYCN to RNA depends on MYCBoxI

To determine whether MYCN directly binds RNA, we recombinantly expressed and purified full-length (FL) MYCN and a series of MYCN deletion mutants (Figure 3A and Figure S4A). We then incubated these constructs with fluorophore-labeled RNA oligonucleotides and measured their binding by gel shift and fluorescence anisotropy assays. We initially used a series of RNA oligonucleotides (62-80 nt) derived from MYCN eCLIP bound intron sequences together with a fragment encompassing the N-terminal region of MYCN (aa 1-154). Fluorescence anisotropy showed that the MYCN N-terminus bound all RNAs with no preference for RNA length (Figure S4B) but with a strong preference for oligonucleotides with higher UG content (Figure 3B). We therefore designed two 80 nt oligonucleotides that differed strongly in UG content (“UG^high^” and “UG^low^”) and performed all subsequent experiments with this pair of oligonucleotides.

**Figure 3:**
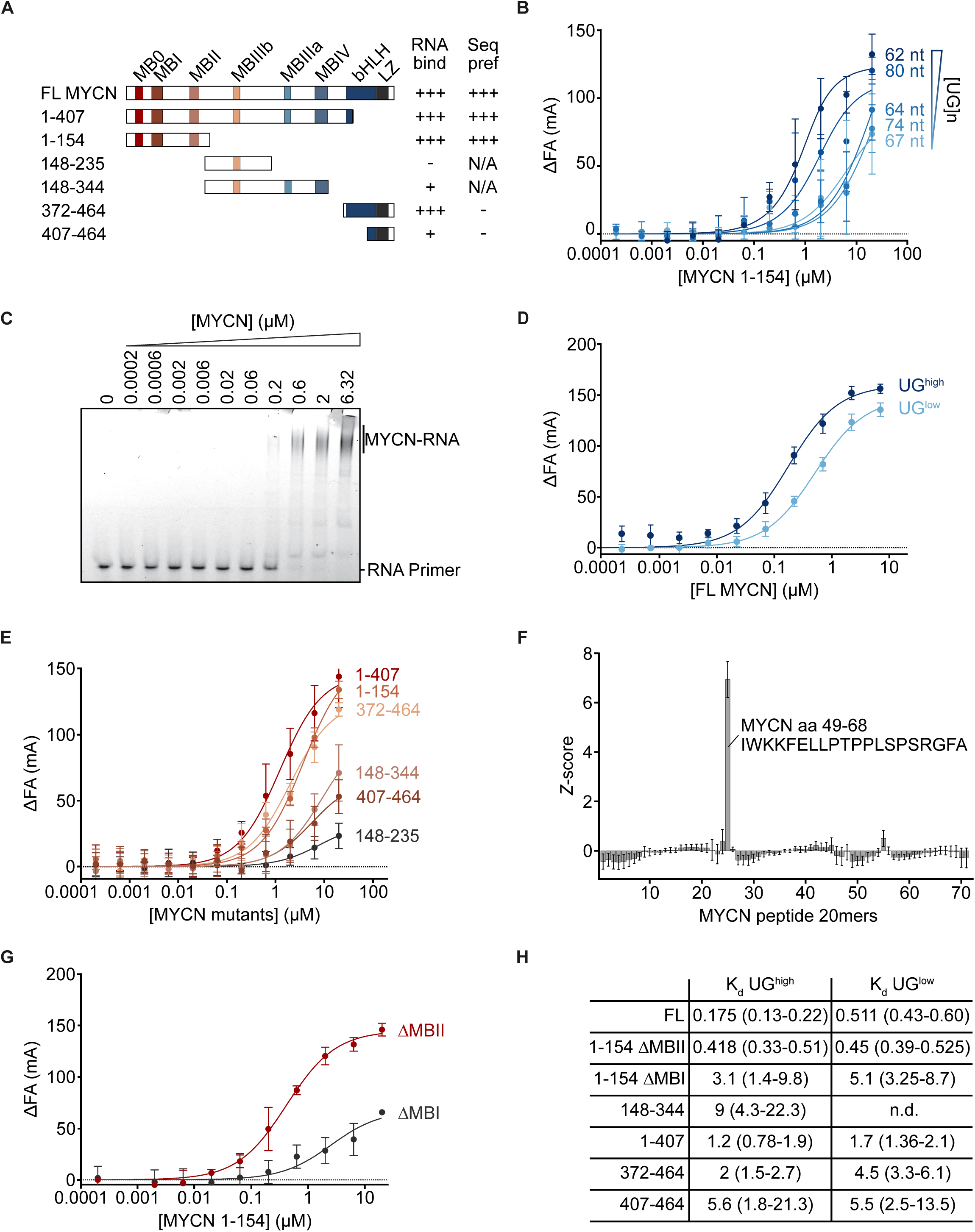
Direct binding of MYCN to RNA depends on MYCBoxI. (**A**) Graphical summary of the purified His-MBP-tagged MYCN constructs that were assayed for RNA-binding by fluorescence anisotropy. (+++) indicates the highest affinity RNA-binding, or highest sequence preference for a UG dinucleotide-rich (UG^high^) compared to UG dinucleotide-poor (UG^low^) RNA oligonucleotide for each construct. MYCN protein domains are labeled on the full-length (FL) MYCN construct diagram. MB: MYC Box. bHLH: Basic Helix-Loop-Helix. LZ: Leucine Zipper. (**B**) Fluorescence anisotropy (FA) measurement using half-log dilutions from 20 μM to 0.2 nM of MYCN 1-154. Five RNA oligonucleotides of varying UG dinucleotide content were used. The relative number of UG-dinucleotides in each construct is indicated on the plot. Dots show mean and error bars depict standard deviation (n=3). (**C**) Electrophoretic mobility shift assay (EMSA) using half-log dilutions from 6.32 μM to 0.2 nM of FL MYCN incubated with 5 nM UG^high^ RNA. The RNA probe was visualized by FITC label (n=3). (**D**) FA measurement using half-log dilutions from 10 μM to 0.2 nM. FL MYCN with UG^high^ or UG^low^ RNA oligonucleotides. Change in anisotropy relative to 5 nM FITC-labeled RNA in the absence of MYCN is plotted relative to FL MYCN concentration. Dots show mean and error bars depict standard deviation (n=3). (**E**) FA measurement of half-log dilutions from 20 μM to 0.2 nM MYCN constructs with UG^high^ RNA oligonucleotide. Change in anisotropy measured as in (D). Dots show mean and error bars depict standard deviation (n=3). (**F**) MYCN aa 1-160 was displayed in microarray format as 72 overlapping oligopeptides (20mers with an overlap of 18 aa and offset of 2). Arrays were incubated with Cy5-labelled RNA 62 nt, which exhibited strongest binding for MYCN 1-154 in (B). Shown are the average *Z*-scores of the measured fluorescence intensities for all tested peptides. Error bars depict standard error of *Z*-score mean. The oligopeptide with the strongest RNA-binding and its sequence are highlighted (n=6). (**G**) FA measurement of half-log dilutions from 20 μM to 0.2 nM MYCN 1-154 mutants lacking MYCBoxI or II with UG^high^ RNA oligonucleotide. Change in anisotropy measured as in (D). Dots show mean and error bars depict standard deviation (n=3). (**H**) Dissociation constant (K_d_) values for the indicated MYCN construct used in FA measurements with UG^high^ and UG^low^ RNA (n=3; n.d.=K_d_ could not be determined).

Both read-outs, gel shift (Figure 3C) and fluorescence anisotropy (Figure 3D), showed that recombinant FL MYCN protein avidly bound the UG^high^ oligonucleotide with a K_d_ value of 0.175 μM and to the UG^low^ oligonucleotide with an affinity of 0.511 μM (see Figure 3H for a summary of K_d_ values). In contrast, MYCN very weakly bound a double-stranded DNA oligonucleotide with the same sequence as the UG^high^ RNA nucleotide (Figure S4C). Testing a series of MYCN deletion mutants of MYCN to determine which regions mediate RNA-binding confirmed that constructs encompassing the N-terminus display the highest affinity for RNA (Figure 3E). While the C-terminal basic region also displays significant affinity for RNA, it did not show any preference for UG-rich sequences (Figure S4D). To map the RNA-binding domains within the MYCN N-terminus, we probed an oligopeptide library spanning the first 160 amino acids of MYCN and consisting of overlapping 20mers with an offset of two amino acids between each peptide with the fluorescent RNA that exhibited the strongest binding for the MYCN N-terminus (62 nt) (Figure 3F). The results showed that a sequence encompassing MYCBoxI, an evolutionary highly conserved domain, exhibited very strong RNA-binding, and that a second sequence, located close to MYCBoxII, which is predicted to bind RNA^34^, may make a minor contribution to RNA-binding. To confirm these data, we expressed MYCN (aa 1-154) proteins lacking either MYCBoxI or MYCBoxII and measured their affinity for the UG^high^ oligonucleotide (Figure 3G). This confirmed that the deletion of MYCBoxI significantly compromised the binding of MYCN to RNA, while a MYCBoxII deletion mutant displayed an affinity similar to FL MYCN (Figure 3H). Collectively, the data show that MYCN binds RNA directly with a preference for UG-rich sequences, recapitulating the association with RNA in cells, and that MYCBoxI is a major RNA-binding domain of MYCN.

### Exosome function controls the dynamics of the MYCN/MAX/MXD network

The data argue that in a cell MYCN exists in DNA– and RNA-bound complexes with different functionalities. Notably, binding of MYCN to RNA reduced its interactions with multiple subunits of the NuA4 histone acetylase complex, arguing that RNA-bound complexes of MYCN do not promote histone acetylation. To understand how binding of MYCN to RNA affects its canonical functions at promoters, we performed spiked ChIP-sequencing experiments (ChIP-Rx) in IMR-32 *MYCN*-amplified cells using a MYCN antibody before and after EXOSC10 depletion. Immunoblots revealed that EXOSC10 loss had no or minor effects on protein levels of MYCN or several proteins that interact with MYCN or MAX at promoters (Figure 4A). In untreated control cells, MYCN associated with most active promoters, consistent with many previous observations (Figures 4B,C). Inspection of multiple individual promoters and global analyses showed that depletion of EXOSC10 led to a significant reduction in MYCN binding to promoters, arguing that the RNA-bound pool of MYCN competes with its promoter-bound pool (Figures 4B,C). As described in the introduction, DNA-bound MYCN forms a heterodimer with MAX; MAX in turn can also form heterodimers with a family of repressor proteins, termed MXD/MNT proteins, which recruit histone deacetylase complexes via a common interaction with the SIN3A adaptor protein. Of the six MXD/MNT proteins, MNT (MXD6) is abundant in neuroblastoma cells^35^ (Figure 4A). We therefore performed CUT&RUN for MAX, MNT, and SIN3A in control and EXOSC10-depleted cells (Figure 4B,C). The results showed that binding of MAX to chromatin also decreased upon EXOSC10 depletion suggesting that MYCN/MAX complexes are the predominant heterodimer present in *MYCN*-amplified neuroblastoma cells. In parallel, binding of MNT and of SIN3A increased at promoters, arguing that MYCN/MAX heterodimers are replaced in part by MNT/MAX heterodimers, which subsequently recruit histone deacetylase complexes (Figure 4B,C). Many effects of MYC oncoproteins occur at virtually all active promoters and are correlated with overall gene expression and transcription^21^. Consistent with these global functions, the extent of both the decrease in MYCN and MAX as well as the increase in MNT/MXD6 and SIN3A paralleled the expression (mRNA) levels of the downstream gene (Figure 4D). The data argue that depletion of EXOSC10 inhibits the transcriptional function of MYCN at promoters. To test this hypothesis, we performed mRNA sequencing from control and EXOSC10-depleted cells. Consistent with this hypothesis, we observed a significant inhibition of MYCN-dependent transcriptional programs, in particular of hallmark MYC-target gene sets involved in protein translation and cell growth; conversely, genes that are repressed by MYCN, for example those involved in the response to interferon alpha, were upregulated upon EXOSC10 depletion (Figure 4E,F). Collectively, the data show that the catalytic subunit EXOSC10 is required to maintain chromatin association of MYCN and prevent replacement of MYCN at promoters with functionally antagonistic and growth-inhibitory members of the MYCN/MAX/MXD network of proteins.

**Figure 4:**
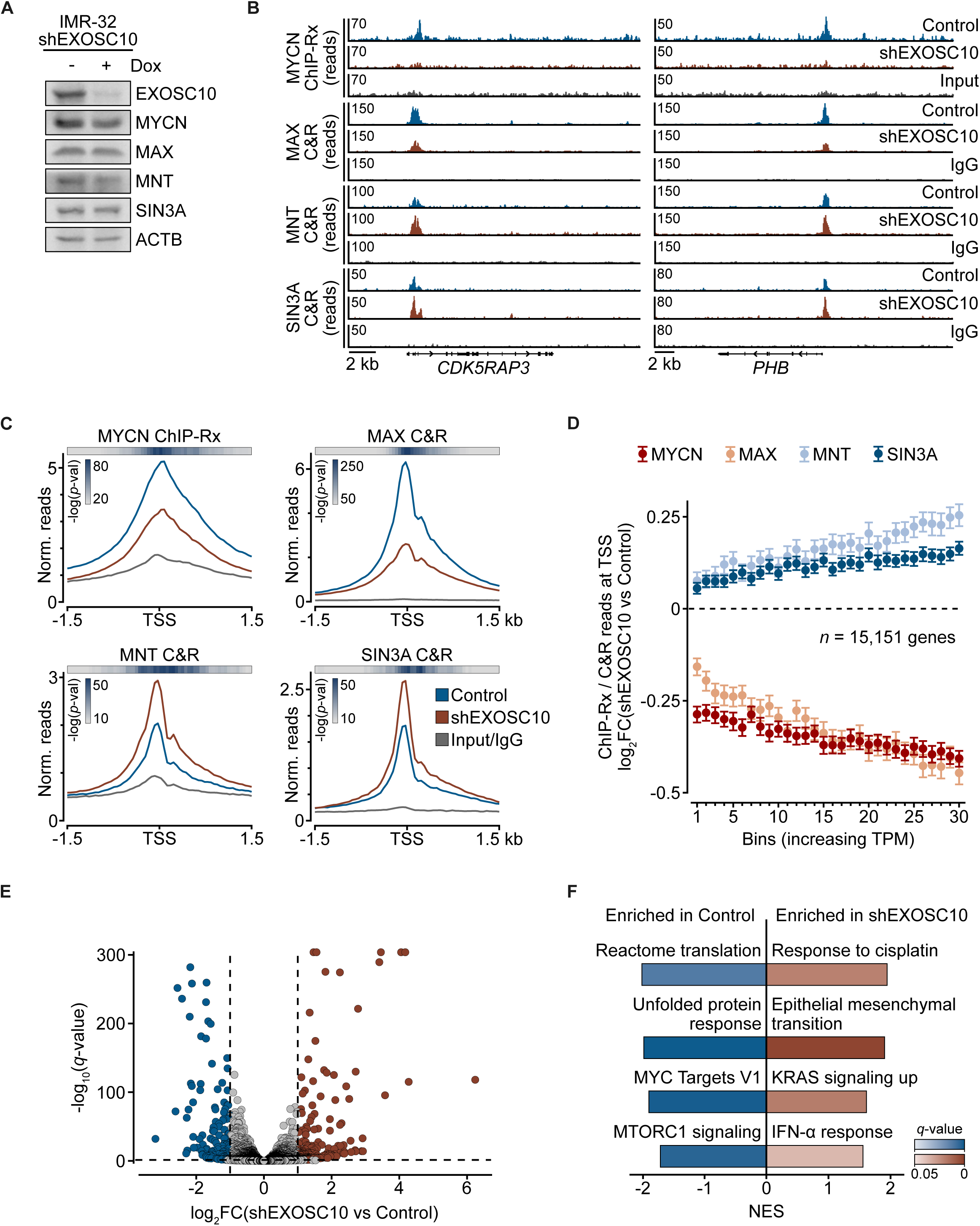
Exosome function controls the dynamics of the MYCN/MAX/MXD network. (**A**) Immunoblot of IMR-32 cells stably expressing an shRNA against EXOSC10 treated for 72 h with doxycycline (Dox) or EtOH as control. Beta actin (ACTB) was used as a loading control (n=3). **(B)** Read distribution of MYCN ChIP-Rx, MAX CUT&RUN (C&R), MNT C&R and SIN3A C&R at the genes *CDK5RAP3* and *PHB*. Cells treated as in (A) (n=3 for MYCN ChIP-Rx; n=2 for all C&Rs). **(C)** Trimmed (2%) average density blots of MYCN ChIP-Rx, MAX, MNT and SIN3A C&Rs centered around the transcription start site (TSS) of all expressed genes (n = 15,151). Cells were treated as in (A). The heatmaps show the *p*-value (two-sided Wilcoxon test) of the difference between the shEXOSC10 and the control condition calculated from the read densities at each individual bin across all genes. (**D**) Correlation between gene expression levels (x-axis) and MYCN, MAX, MNT or SIN3A change in occupancy at TSS (y-axis) following EXOSC10 depletion. Expressed genes (n = 15,151) were grouped into 30 bins (505 genes per bin) of increasing TPM (transcripts per million mapped reads) (n=3). Points are the mean of log_2_FC between shEXOSC10 and control condition and the error bars depict 95% confidence intervals (n=3). **(E)** Volcano plot of mRNA sequencing data showing differences in mRNA levels upon EXOSC10 loss. Cells were treated as in (A). The x-axis shows the log_2_FC between EXOSC10 depleted and control cells and the y-axis depicts statistical significance (*q*-value). Genes were considered significantly altered if passing *q*-value < 0.05 and log_2_FC > 1 or < –1 thresholds (n=3). **(F)** Gene set enrichment analysis of mRNA sequencing data as described in (E). Shown is a selection of significantly up– and downregulated gene sets. NES: Normalized enrichment score.

### MYCN associates with the NEXT targeting complex of the exosome

To identify possible effector functions of MYCN when bound to RNA, we first sought to better understand its interactions with the exosome. The nuclear exosome does not recognize target RNAs directly, but is engaged by specific targeting complexes that recognize different RNA species^27^. To investigate whether MYCN associates with any known targeting complex, we performed immunoprecipitation experiments from lysates of *MYCN*-amplified neuroblastoma cells. To test the dependence of the interaction on the exosome, we used IMR-32 EXOSC5 AID cells before and after the addition of auxin (IAA) (Figure 5A). As a positive control, we probed MYCN immunoprecipitates for three MYCN cofactors involved in transcriptional regulation, MAX, TRRAP and the CDC73 subunit of the PAF1 complex, and confirmed their interaction with MYCN. We also confirmed that EXOSC5 co-immunoprecipitates with MYCN, as reported previously (Figure 5A)^24,36^. Intriguingly, ZCCHC8 and RBM7, which are two specific subunits of the <u>n</u>uclear <u>ex</u>osome targeting (NEXT) complex^37,38^ that targets the exosome mainly to non-polyadenylated RNAPII-derived RNAs, including pre-mRNAs and PROMPTs, efficiently co-immunoprecipitated with MYCN (Figure 5A). Conversely, PABPN1 and ZFC3H1, two specific subunits of the <u>p</u>oly(<u>A</u>) tail e<u>x</u>osome targeting (PAXT) complex, which targets the exosome to polyadenylated transcripts, did not co-immunoprecipitate with MYCN^36^; MTR4, a helicase present in both complexes, co-immunoprecipitated with MYCN with intermediate efficiency (Figure 5A). Conversely, ZCCHC8, but not PABPN1 immunoprecipitates, showed a robust interaction with MYCN (Figure S5A).

**Figure 5:**
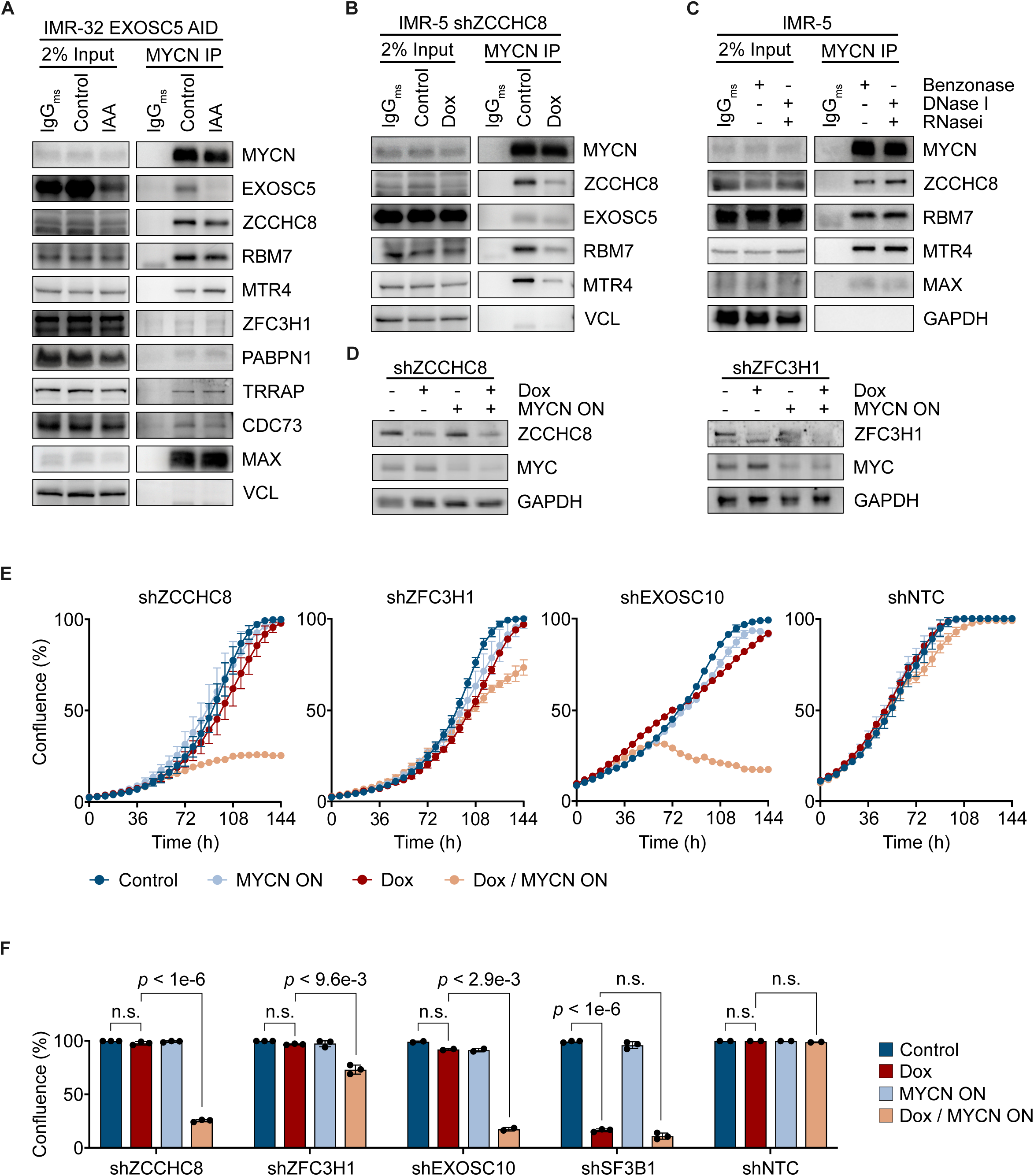
MYCN associates with the NEXT targeting complex of the exosome. (**A**) Immunoblot of endogenous MYCN immunoprecipitations (IPs) from IMR-32 EXOSC5 AID cells treated for 1.5 h with IAA (100 µM) or DMSO as control. Samples were treated with 50 U/ml benzonase before precipitation. Vinculin (VCL) was used as loading control. Higher exposure time for α-TRRAP and α-CDC73 in IP panel compared to corresponding input, samples were on the same membrane (n=2). **(B)** Immunoblot of endogenous MYCN IPs from IMR-5 cells that express a doxycycline-inducible shRNA targeting ZCCHC8. Cells were treated for 48 h with 1 µg/ml Dox (control with EtOH). Vinculin (VCL) was used as loading control (n=2). (**C**) Immunoblot of endogenous MYCN IPs from IMR-5 cells. Samples were treated with benzonase or DNAse in the presence of RNAse inhibitor where indicated (each 50 U/ml for 1 h). GAPDH was used as loading control (n=1). **(D)** Immunoblot of SH-EP MYCNER cells expressing Dox-inducible shRNAs targeting ZCCHC8 (left) or ZFC3H1 (right). Cells were treated for 48 h with 1 µg/ml Dox and 4 h with 100 nM 4-OHT (MYCN ON), where indicated. MYC was used as a control of MYCNER activation, as 4-OHT addition will decrease MYC protein levels. GAPDH was used as loading control (n = 2). **(E)** Growth curves of SH-EP MYCNER cells stably expressing shRNAs targeting ZCCHC8, ZFC3H1, EXOSC10, or a non-target control (NTC). Cells were treated with 1 µg/ml Dox and 100 nM 4-OHT (MYCN ON) as indicated over the entire measurement time. Dots show mean and error bars depict standard deviation (n=3 for ZCCHC8, ZFC3H1 and SF3B1; n=2 for EXOSC10 and NTC). (**F**) Endpoint confluency (t = 144 h) from growth curves in (E) including SH-EP MYCNER cells expressing SF3B1 shRNA. Dots show mean and error bars depict standard deviation. *P*-values were calculated with a multiple paired Welch’s t-test; n.s. = not significant.

The interaction of MYCN with NEXT complex subunits was unaffected by auxin dependent degradation of EXOSC5, arguing that it does not depend on an intact exosome (Figure 5A). To determine whether MYCN interacts with an intact NEXT complex, we generated IMR-5 cells that express a doxycycline-inducible shRNA targeting ZCCHC8, a core subunit of the NEXT complex^37,39^. Depletion of ZCCHC8 not only reduced its interaction with MYCN, but also the interaction of MYCN with MTR4 and RBM7, demonstrating that MYCN interacts with the intact NEXT complex (Figure 5B). MYCN interactions with all three NEXT subunits did not decrease when precipitations were performed after digestion of both DNA and RNA by benzonase, and conversely, did not increase when RNAs were preserved after DNase treatment by an RNAse inhibitor (Figure 5C), suggesting that the interaction with MYCN does not depend on RNA contained in the complex.

The growth of *MYCN*-amplified cells depends on exosome function, whereas the growth of neuroblastoma cells expressing MYC is much less dependent on the exosome^24^. To test whether this extends to the NEXT complex, we expressed shRNAs targeting ZCCHC8 in SH-EP-MYCNER neuroblastoma cells; SH-EP cells normally express MYC, but addition of 4-hydroxytamoxifen (4-OHT) activates MYCN (MYCN ON), which in turn represses MYC expression^40^ (Figure 5D). Depletion of ZCCHC8, like that of EXOSC10, strongly suppressed the growth of cells in the presence of 4-OHT, but had little effect on cell growth in the absence of 4-OHT (Figures 5E,F). In contrast, an shRNA targeting the ZFC3H1 subunit of the PAXT complex did not slow the proliferation of these cells, despite strong depletion of ZFC3H1 (Figures 5D-F). Depletion of SF3B1, an essential splicing factor, suppressed growth under both conditions, as expected (Figure 5F and Figure S5B). We concluded that a significant fraction of the NEXT complex associates with MYCN in *MYCN*-amplified neuroblastoma cells and that this complex is essential for their growth.

### RNA-bound MYCN promotes the degradation of non-polyadenylated intronic RNA

Targeting complexes like NEXT direct the nuclear exosome to specific classes of RNAs, raising the possibility that MYCN affects RNA recognition by the exosome in neuroblastoma cells. To determine the impact of MYCN on the exosome RNA-binding, we expressed a catalytically inactive allele of EXOSC10 (EXOSC10_mut_) in IMR-32 cells. This construct carries D313A and E315A mutations in a conserved 309-EFAVDLEHHS-318 motif that is required for RNA degradation ^41^. We have shown previously that ectopic expression of EXOSC10_mut_ increased the number of PROMPTs, consistent with published data that the RNA exosome degrades PROMPTs^24^ and immunoblots established that this allele interacts with the core exosome subunit, EXOSC5 (Figure S5C). To probe for the effect of MYCN, we co-expressed two doxycycline-inducible shRNAs that target MYCN (Figure 6A) and performed eCLIP experiments for EXOSC10_mut_ in both control and MYCN depleted conditions (Figure 6B). We performed stringent peak calling as before and then inspected the EXOSC10_mut_ read distribution around the called high-confidence eCLIP peaks for both conditions (Figures 6B,C). In control and MYCN depleted cells, density plots and heatmaps centered around the respective eCLIP peaks showed a sharp peak of EXOSC10_mut_ RNA-binding surrounded by a broad region of further binding, which can extend for several kilobases (Figures 6C,D). We identified 538 and 489 high-confidence peaks in control and MYCN depleted cells, respectively (Figure 6E). While EXOSC10_mut_ was predominantly bound to introns in both control and MYCN-depleted conditions (Figure 6E), depletion of MYCN caused a significant redistribution of the exosome on intronic RNA. Specifically, binding sites bound by EXOSC10_mut_ in the presence of MYCN were more strongly localized in 5’ proximal introns, mirroring RNA-binding of MYCN itself (Figure 6F). To investigate whether MYCN and EXOSC10_mut_ bind similar RNA sites, we plotted MYCN eCLIP reads around the high-confidence EXOSC10_mut_ eCLIP peaks with and without MYCN depletion (Figure 6G). We observed that MYCN co-localizes strongly with EXOSC10_mut_ control peaks, but this co-localization is blunted upon MYCN depletion indicating that MYCN re-localizes the exosome from sites with no or little MYCN binding to sites that overlap with MYCN binding sites (Figure 6G and Figure S5D). Taken together, the findings argue that MYCN enhances association of the exosome with RNA close to the former’s binding sites.

**Figure 6:**
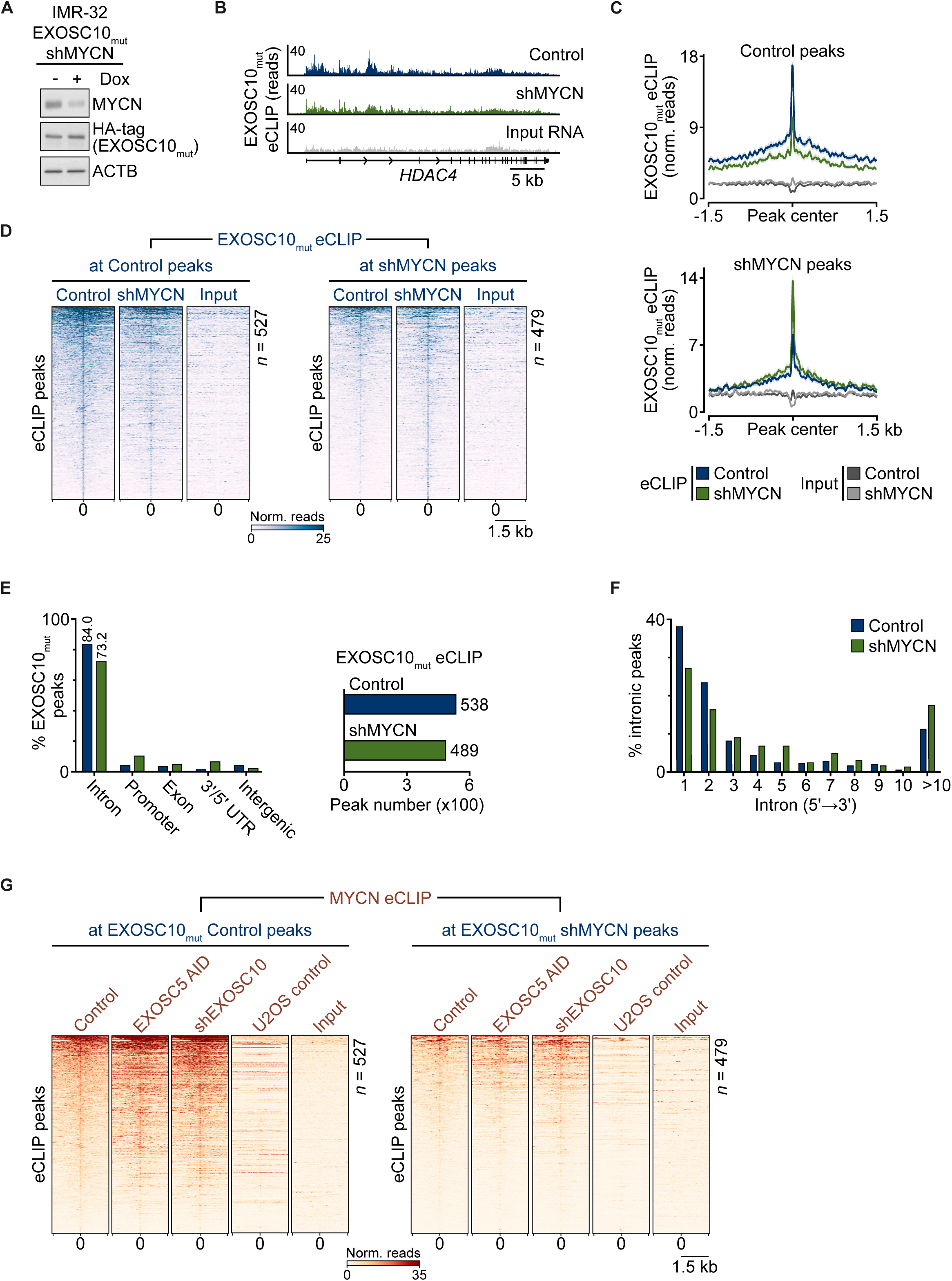
MYCN enhances RNA binding of the nuclear exosome at its binding sites. (**A**) Immunoblot of IMR-32 cells co-expressing HA-tagged EXOSC10 D313A/E315A inactive mutant (EXOSC10_mut_) and two doxycycline (Dox)-inducible shRNAs against MYCN. Cells were treated for 72 h with Dox (1 µg/ml) or EtOH as control. ACTB was used as a loading control (n=3). (**B**) Read distribution of EXOSC10_mut_ eCLIPs at the *HDAC4* gene. Cells treated as in (A). Input RNA corresponds to the control condition size-matched eCLIP input (n=2). (**C**) Trimmed (2%) average density blots of EXOSC10_mut_ eCLIP and corresponding size-matched input reads in depicted conditions, centered around EXOSC10_mut_ control (left; n=527) and MYCN-depleted (right; n=479) high-confidence eCLIP peaks. Only reads in the sense strand are shown. Lines correspond to mean of biological duplicates, shaded regions depict standard error. (**D**) Heatmaps of trimmed (2%) reads of EXOSC10_mut_ eCLIPs centered around EXOSC10_mut_ control or shMYCN high-confidence eCLIP peaks. Input corresponds to the RNA input of the EXOSC10_mut_ eCLIP control condition. Heatmaps were sorted based on descending mean reads in the EXOSC10_mut_ eCLIP control condition. Only reads in the sense strand are shown. Shown are the means of biological duplicates. (**E**) Peak distribution at depicted gene regions for EXOSC10_mut_ eCLIP control and MYCN-depleted condition. (**F**) Distribution of intronic EXOSC10_mut_ eCLIP peaks from nearest to farthest from the transcription start site. (**G**) Heatmaps of trimmed (2%) reads of MYCN eCLIPs centered around EXOSC10_mut_ control or shMYCN high-confidence eCLIP peaks. Depicted input corresponds to the MYCN eCLIP control condition. All heatmaps were sorted based on descending mean reads in the MYCN eCLIP control condition. Only reads in the sense strand are shown. Shown are the means of biological duplicates.

To determine the function of MYCN’s interaction with EXOSC10, we used IMR-32 cells stably expressing doxycycline-inducible shRNAs targeting either EXOSC10 or MYCN and mapped the 3’ ends of nascent, 4sU-labelled RNAs (3’RNAseq). To do so, we used oligo-dT primers to prime reverse transcription either directly (mapping polyadenylated RNAs – “only polyA”) or after incubation of nascent RNA with a bacterial poly(A)-polymerase (all 4sU-labelled RNAs will be isolated irrespective of *in cellulo* polyadenylation – “all RNAs”)^42^. In line with its described function, EXOSC10 depletion significantly increased RNA 3’ ends downstream annotated transcription start sites of PROMPTs, however MYCN loss had no discernible effect at those positions (Figure S6A). We then turned to the effects of EXOSC10 and MYCN loss at introns, as both MYCN and EXOSC10_mut_ bind long introns (> 10k nt in all conditions), which are known to be trimmed by the exosome^42^ (Figure S6B). We plotted the ratio at MYCN bound introns between untreated control cells and cells depleted of EXOSC10 or MYCN and demarcated the effects between normally polyadenylated RNAs and RNAs that additionally include non-polyadenylated RNAs. EXOSC10 loss increased the number of 3’ ends at all MYCN-bound intron-exon boundaries (Figures 7A,B and Figure S6C). Depletion of MYCN had no effect on polyadenylated 3’ ends, but significantly increased the number of non-polyadenylated 3’ ends at its bound introns, arguing that MYCN promotes degradation of non-polyadenylated target introns (Figures 7B,C and Figure S6C). Inspection of 3’ ends at all IMR-32 expressed introns showed that the effects of EXOSC10 and MYCN depletion are not global, but rather that the MYCN bound introns represent a subset of exosome-targeted introns (Figure 7B, bottom). Specifically, stratification for intron length demonstrated that MYCN depletion effects are restricted to long introns (>4kb) (Figure 7C and Figure S6D). Taken together, the data show that MYCN promotes the degradation of long introns to which it is bound via the nuclear exosome. A model summarizing the findings presented in this work is shown in Figure 7D.

**Figure 7:**
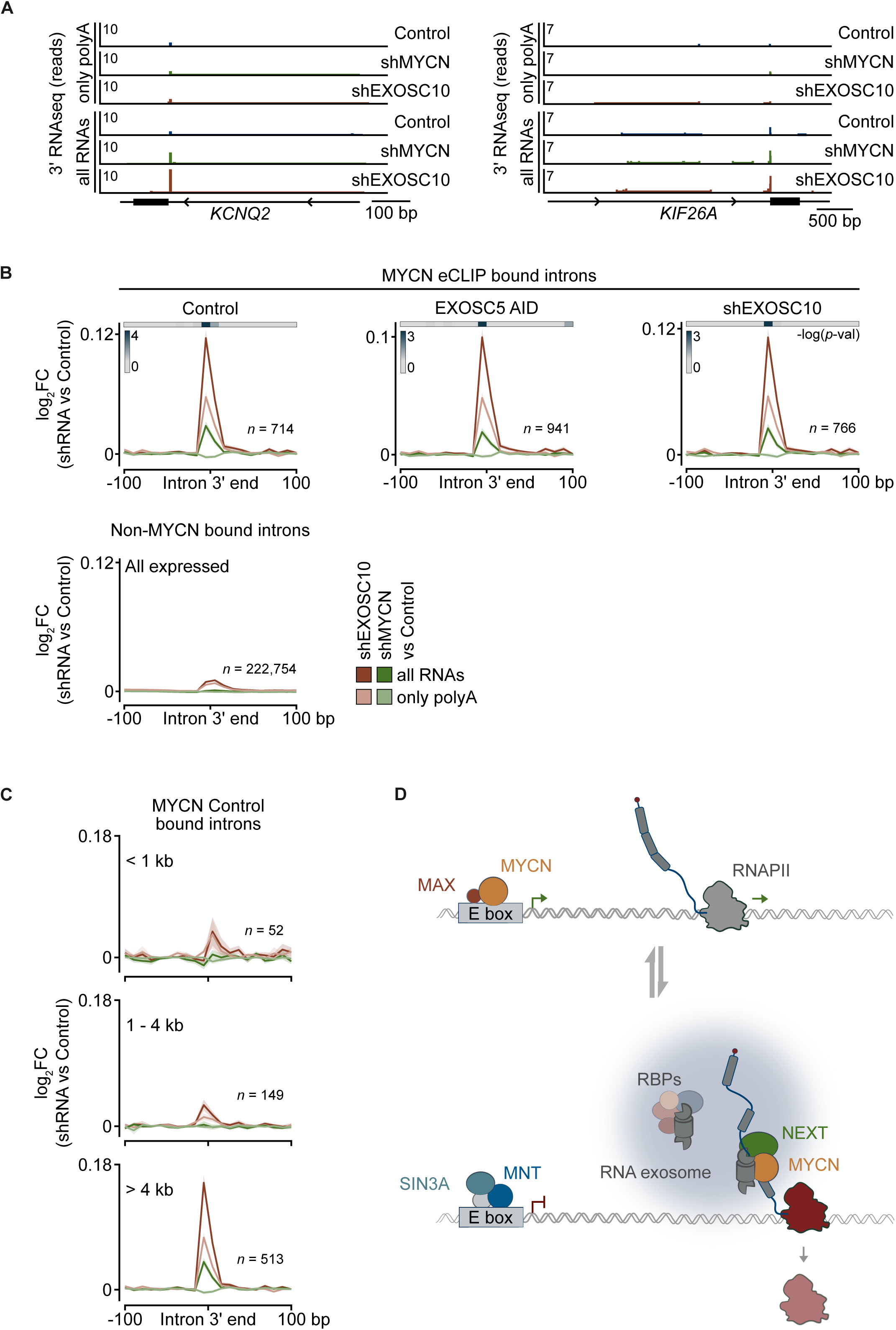
RNA-bound MYCN promotes the degradation of non-polyadenylated intronic RNAs. (**A**) Read distribution of 3’ RNAseq reads at the intron-exon boundaries (intron 3’ end) of the *KCNQ2* and *KIF26A* genes. IMR-32 shEXOSC10 or shMYCN cells were treated with Dox (1 µg/ml) for 72 h. Control condition treated with EtOH. RNA libraries were generated with (all RNAs) or without (only polyA) an RNA polyadenylation step to also isolate non-polyadenylated RNA 3’ ends (n=2). (**B**) Log_2_FC between EXOSC10 or MYCN depletion compared to control condition at the 3’ ends of introns bound by MYCN in the depicted eCLIP conditions. Samples containing polyadenylated and non-polyadenylated RNAs (all RNAs; darker colors), and only polyadenylated RNA (only polyA; lighter colors) are shown (top). The same analysis is shown for all introns of IMR-32 expressed genes (bottom). Only reads in the sense strand are shown. Lines correspond to mean of biological duplicates, shaded regions depict standard error. (**C**) Log_2_FC between EXOSC10 or MYCN depletion compared to control condition at the 3’ ends of introns bound by MYCN in eCLIP untreated control cells. Introns are stratified according to length as shown. (**D**) Model summarizing our findings.

## Discussion

The MYCN oncoprotein, like its counterpart MYC, forms a heterodimeric DNA-binding complex with MAX and regulates the transcription of a broad spectrum of target genes. Two evolutionary conserved domains of MYC proteins are critical for this function^12^: the C-terminal BR-HLH-LZ domain mediates heterodimerization with MAX and binding to DNA, and a short domain termed MYCBoxII recruits TRRAP, a scaffold protein of several histone acetylase complexes (see Introduction). Our previous work had shown that MYCN is associated with the nuclear exosome and that MYCN-driven tumor cells depend on exosome function for passage through the S phase of the cell cycle^24,26^. This was surprising because the exosome is a 3’-5’ RNA exonuclease and is not known to be involved in transcriptional regulation. We now show that the MYCN/MAX heterodimer represents only one of two distinct physical states of MYCN. In its second state, MYCN forms stable high molecular weight complexes containing the nuclear exosome and other RNA-binding proteins. The shift between the two states occurs during the transition from G1 to S phase. The assembly of high molecular weight MYCN complexes can also be triggered by loss of the EXOSC10 catalytic subunit of the exosome, which is present in a fraction of the high molecular weight MYCN complexes. The data show that MYCN either stably associates with nascent RNA in these complexes, or that association with RNA triggers their assembly, for example by altering MYCN conformation or by blocking MYCN turnover by the proteasome (see below).

Consistent with these observations, MYCN binds to thousands of sites on RNA, mainly localized in introns, arguing that MYCN associates with unprocessed mRNAs. The eCLIP method that we used maps direct RNA-protein contacts after UV crosslinking, which however preferentially crosslinks RBPs to uridines and guanosines^31,43^. Together with the recombinant MYCN protein binding assays we conclude that MYCN directly binds UG-rich RNA sequences with high affinity and preclude the possibility that the eCLIP bound UG-sequence motifs are an artifact of UV crosslinking. Two regions in MYCN, MYCBoxI and the basic region have significant affinity to RNA, with MYCBoxI showing a clear preference for UG-rich sequences while the carboxy-terminal basic region has no detectable sequence specificity. MYCBoxI is highly conserved among all MYC proteins and contains a degron that is recognized by the FBXW7 ubiquitin ligase and the USP28 deubiquitylating enzyme^44,45^; in MYCN, MYCBoxI also harbors the binding site of Aurora-A, which antagonizes proteasomal degradation by FBXW7^46,47^. Moreover, MYCBoxI also contains three conserved phosphosites that in MYCN are targeted by GSK3 (for T58)^48^, CDK1 (for S62)^49^ and an unknown kinase (for S64). Taken together, these data argue that RNA-binding of MYCN and its complex formation with RNA-binding proteins are tightly regulated processes and that other MYC family members may also bind RNA. The latter hypothesis is supported by several recent observations^50–52^.

Proteomic analyses showed that RNA-bound complexes of MYCN are functionally distinct from their DNA-bound counterparts. Specifically, RNA-associated complexes of MYCN did not contain the major MYCN-associated histone acetylase complex (NuA4), suggesting that they do not promote histone acetylation. Furthermore, exosome loss increased the amount of MYCN bound to RNA but decreased promoter binding, arguing that RNA– and DNA-binding compete for the available pool of MYCN molecules. Conversely, promoter binding of MNT, a repressive partner of MAX, and of SIN3A, an adaptor protein via which MNT and MXD recruit histone deacetylase complexes, increased globally, corresponding with a downregulation of the MYCN-dependent gene expression program. The data argue that the MYCN/MAX/MXD network globally adapts the chromatin state in response to the completion of mRNA processing.

Providing a feedback signal to the MYC/MAX/MXD network is not the sole function of RNA-bound MYCN complexes. To recognize its substrates, the exosome relies on different targeting complexes and MYCN is predominantly associated with the NEXT complex, and to a significantly lesser degree with the PAXT complex. This association has functional relevance, as eCLIP experiments showed that MYCN enhances exosome binding to hundreds of intronic sites, and RNA 3’ end mapping demonstrated that MYCN promotes degradation of non-polyadenylated mRNAs at its bound introns. The data thus argue that MYCN functions as an accessory factor of NEXT and the exosome. Notably, the 3’ ends of long introns, which are predominantly bound both by MYCN and EXOSC10, have been shown to be primary targets of the exosome by both RNAi depletion and auxin induced degradation experiments^42^, arguing that also the function of MYCN in trimming the 3’ ends of long introns is a primary effect and not a by-product of exhausted 5’-3’ exonucleases.

The results are compatible with several non-mutually exclusive models. For example, MYCN may act co-transcriptionally in a pathway leading to termination of inefficient RNAPII transcription^24^. Supporting this model is the observation that MYCN binding sites are not restricted to splice sites but often broadly distributed along the lengths of introns. RNAPII transcription is characterized by frequent pauses during early elongation, which could explain MYCN’s preferential binding to 5’ proximal introns of long genes^53^. Furthermore, RNAPII frequently encounters the replication fork during S phase^54^, which may account for the shift of MYCN from MYCN/MAX to RNA-bound complexes during this cell cycle phase. Stimulation of intronic RNA degradation by the exosome may also be a mechanism by which MYCN limits R-loop formation^40^. Either of these processes is expected to enhance the resilience of neuroblastoma cells to stresses arising from a stalling RNAPII, for instance by preventing conflicts of the replication and transcription machineries. Ultimately, our findings suggest that the ability to control exosome functions at introns may significantly contribute to the oncogenic function of MYCN. The dependence on the exosome and the NEXT complex is particularly strong for MYCN-driven tumor cells. Since MYC and MYCN are expressed in a mutually exclusive manner in tumors, expression of MYCN may identify cancer cells with a specific and targetable vulnerability. We therefore propose that targeting the function of MYCN in exosome-driven RNA turnover may open a significant therapeutic window for *MYCN*-amplified neuroblastoma and other MYCN-driven tumors.

## Acknowledgements

This work was supported by grants from the European Research Council (SENATR ERC #101096948 to M.E.), the German Cancer Aid via the Excellence Program (#70114538 to M.E.), the Mildred Scheel Junior Research Center Program (to D.P. and G.B.), the German Research Foundation (EI 222/21-1 and INST 93/1023-1-FUGG to M.E.) and the Alex’s Lemonade Foundation Crazy 8 Initiative (to S.M.V. and M.E.). The authors thank Carsten Ade, Barbara Bauer, Ulrike Samfaß and Tobias Roth for technical support, Cornelius Schneider for initial help with gradients, Raphael Silveira Vidal for bioinformatics advice and Theresa Hauck for preparing MYCN deletion constructs.

## Author Contributions

Conceptualization, D.P., S.M.V. and M.E.; Methodology, D.P., S.A.H., D.F., L.U. and M.E.; Investigation, D.P., S.A.H., D.F., L.U., K.S., I.M., T.J.R., A.B., O.R.V. and C.S.-V.; Formal Analysis, D.P., I.M. and P.G.; Visualization, D.P., S.A.H, D.F., L.U., O.R.V. and T.J.R.; Writing – Original Draft, D.P. and M.E.; Supervision, P.B., G.B., S.M.V, H.M.M. and M.E.; Funding Acquisition, D.P., S.M.V., G.B. and M.E.

## Declaration of interests

M.E. is a founder and shareholder of Tucana Biosciences.

## STAR Methods

### Cell culture

Human IMR-32 (male), IMR-5 (male) and SH-EP (female), HEK293TN (female) and murine NIH3T3 (not applicable) cell lines were cultured at 37°C and 5% CO_2_. IMR-32, IMR-5 and SH-EP cells were cultured in RPMI 1640 medium (Thermo Fisher Scientific), while the HEK293TN and NIH3T3 were cultured in DMEM (Thermo Fisher Scientific) medium. All media were supplemented with 10% fetal bovine serum (FBS; Capricorn Scientific) and 1% penicillin/streptomycin (P/S; Sigma-Aldrich). Cell lines were validated by STR analysis and regularly tested for mycoplasma contamination. Cell lines containing puromycin resistance constructs were regularly re-selected via incubation in 1 µg/ml puromycin-containing medium for 72 h. Cells were synchronized at the G1/S boundary by incubating in 2 mM thymidine-containing medium (Sigma-Aldrich) for 16 h, releasing by washing with PBS and incubating in normal medium for 8 h, and then re-blocking in 2 mM thymidine-containing medium for another 16 h. The cells were then washed with PBS and cultured in normal medium before being harvested at 4 h (S phase), 8 h (G2/M phase) and 13 h (G1 phase). For live growth imaging, cells were seeded in 48-well plates (Greiner). Treatment was started 12 h after seeding and treatment medium was refreshed every 48 h. Cells were incubated for 6 days and phase-contrast confluence was monitored by time-lapse microscopy using the Incucyte SX5 system, with 4 fields analyzed every 6 h using 10x magnification. All fields contained cells at the start of measurements.

### Flow cytometry analysis (FACS)

For propidium iodide (PI) FACS, cells were harvested by trypsinization, washed with cold PBS and then fixed in 80% v/v ethanol overnight at –20°C. After washing with PBS, the pellets were resuspended in sodium citrate (38 mM) containing PI (54 µM) and RNase A (24 µg/ml) and incubated for 30 min at 37°C in the dark. Subsequently, the samples were analyzed on a BD FACSCanto II flow cytometer (BD Sciences) using the BD FACSDIVA software.

### General cloning

The EXOSC5 auxin-inducible degron (AID) construct was cloned as in ^56^. To generate the pJET-NAID-Blast-entry vector (N-terminal tag), a gBlock (IDT) with the AID sequence was cloned into pJET1.2. The homology-directed repair (HDR) template was obtained by PCR-amplifying homology arms (HA) using genomic DNA from isolated IMR-32 cells as template. The PCR fragments were digested with AgeI and MluI, or EcoRI and SpeI enzyme pairs, respectively. The HAs were then cloned into the entry vector, resulting in pJET-EXOSC5-HDR. The guide RNAs sgR1 and sgR2 were cloned into PX458 vector (Addgene: 48138). The plasmid containing the TIR1 was already available in the lab.

For RNA binding assays, the gene for full-length MYCN (1-464) was amplified from cDNA and inserted behind a 6xHis-Maltose binding protein (MBP)-Tobacco etch virus cleavage site via ligation independent cloning (Vector 1C, Addgene: 29654). Truncation mutants were amplified from the full-length gene and inserted in the 1C vector. The MycBoxI (deletion of residues 45-65) and MycBoxII (deletion of residues 111-134) constructs were synthesized as gBlocks and the sequence was optimized for E. coli expression (IDT). The full-length MAX gene was synthesized by Twist Bioscience and was inserted into the 1C vector via ligation independent cloning. All of the constructs described in this section were verified by Sanger Sequencing.

The shRNA-mirE sequences for MYCN, EXOSC10, ZCCHC8, ZFC3H1, and SF3B1 were selected using a modified SplashRNA algorithm^57^ and DNA oligos were ordered from Sigma. After PCR amplification with primers containing XhoI and EcoRI restriction sites the digested constructs were inserted into the vector pLT3GGmirEPPIR^58^ containing a PGK promoter and used to transform XL-1 blue *E.coli* cells.

### Generation of auxin inducible degron (AID) knock-in cell line

Cells were cultured in 10 cm culture plates and transfected with pJET-EXOSC5_HDR and PX458_EXOSC5_sgR1 or PX458_EXOSC5_sgR2 plasmids using PEI as the transfection reagent. Untransfected cells were used as negative selection controls. Cells were trypsinized 72 h after transfection and seeded into five 15 cm plates (serial dilutions of 1:5, 1:10, 1:50, 1:100 and the remaining cells). Cell were selected by adding 1 mg/ml Blasticidin (InvivoGen) and replacing with fresh antibiotic-containing medium every 48 h. When the negative selection control cells were completely dead, the transfected cells were cultured in fresh medium without antibiotics. Individual clones were then picked and transferred to 24-well plates. Genomic PCR and immunoblotting were used to identify homozygous clones. For TIR1 expression, lentivirus containing the plasmid pRRL-SFFV-OsTir1_3x_MYC_tag_T2A-eBFP2 was produced in HEK293T cells. The homozygous EXOSC5 AID clones were infected with the virus-containing supernatant and sorted for BFP expression by FACS analysis 48 h after transduction. EXOSC5 AID degradation was validated by immunoblotting after treatment with 100 µM indole-3-acetic acid (IAA) at various time points.

### Purification of MYCN constructs

MYCN constructs were expressed in BL21 (DE3) RIL LOBSTR cells. Briefly, overnight cultures were used to inoculate 4-6 L of 2xYT media. Cells were grown at 37°C, shaking in baffled flasks until reaching an OD 600 of 0.4-0.6. The temperature was then lowered to 18°C and protein expression was induced by the addition of 0.5 mM IPTG. Cells were grown for an additional 16 hrs. Cells were collected by centrifugation, resuspended in lysis buffer (500 mM NaCl, 20 mM Tris-HCl pH 7.9 at 25°C, 10% glycerol, 30 mM imidazole pH 8.0, 5 mM beta-mercaptoethanol, 0.284 µg ml−1leupeptin, 1.37 µg ml−1 pepstatin A, 0.17 mg ml−1 PMSF, and 0.33 mg ml−1 benzamidine). Resuspended cells were either used directly for protein purification or snap frozen in liquid nitrogen. All protein purification steps were performed at 4°C unless otherwise noted. Cells were lysed by sonication and lysates were clarified by centrifugation. Clarified lysate was applied to a 5 mL HisTrap column equilibrated in lysis buffer. The column was washed with lysis buffer and additionally washed with a high salt buffer (lysis buffer with 1 M NaCl) for 5 CV before returning to lysis buffer. Protein was eluted from the HisTrap column with nickel elution buffer (200-500 mM NaCl, 20 mM Tris-HCl pH 7.9, 500 mM imidazole pH 8.0, 10% glycerol, and 5 mM BME) onto a 10 mL amylose column (New England Biolabs). The amylose column was washed with lysis buffer. Protein was eluted from the amylose column in amylose elution buffer (200-500 mM NaCl, 20 mM Tris-HCl pH 7.9, 30 mM imidazole pH 8.0, 116 mM maltose, 10% glycerol, and 5 mM BME). Protein purity was assessed by SDS-PAGE followed by Coomassie staining. Fractions containing MYCN were pooled and concentrated with an Amicon Millipore ultrafiltration device (10000 MWCO). Concentrated protein was applied to a HiLoad Superdex 200 16/600pg column (GE) equilibrated in size exclusion buffer (200-500 mM NaCl, 20 mM Tris-HCl pH 7.9, 10% glycerol, and 1 mM DTT). The FL MYCN construct was applied to a Superose 6 10/300 size exclusion column (GE). Protein purity was assessed by SDS-PAGE followed by Coomassie staining. Fractions containing MYCN were pooled and concentrated with an Amicon Millipore ultrafiltration device (10000 MWCO). Protein concentration was determined using the calculated extinction coefficient for His6-MBP-MYCN and the measured absorbance at 280 nm. Protein was aliquoted, snap frozen in liquid nitrogen, and stored at –80°C until use.

### Virus production and transduction

HEK293TN cells at 70-80% confluence were transfected with construct-containing plasmids, along with the packaging plasmids psPAX2 and pMD2G, using polyethylenimine (PEI; Sigma-Aldrich) in transfection medium (2% FBS, no P/S). The transfection medium was replaced with normal medium 6 h after transfection and virus-containing supernatant was collected 48 and 72 h after transfection using 0.45 µm filters. Cells were transduced with polybrene and selected 48 h after transduction using 0.5-3 µg/ml puromycin for at least 72 h or by FACS sorting for GFP-, RFP– or BFP-positive cells.

### Fluorescence anisotropy assays for RNA and DNA binding

5’ 6-FAM labelled RNAs were purchased from IDT with the following sequences: “62” 5’-6-FAM-UGG GCC UGG AAU AGU GGG GGC CCA GGG GAA GCU GCG AUG UGC UUC UGA GUG UUU GGA ACG AG-3’, “64” 5’-6-FAM-GAG AUG GCG CAU GGG CCA GGC UAG ACC AGC CGC UGG CCC AGA CGC CCU UUC UGC CUA AAC AAC C-3’, “67” 5’-6-FAM-CUU UCC CAG AGG UUA GCA AGG CAU GAC AGC CAC CGC CAU CAU CAU UGG CAU CAU CAC CAC CAU CAU C-3’, “74” 5’-6-FAM-UUU UCC CGA CUG GCU GAG AUU CCC GCA GCA UCC CGA CUC CGC UUC AGG GCU CCU GUG AGG ACA GGG ACU AUG GC-3’, “80” 5’-6-FAM-UUG GGG UGA GUG UGA CUU CUG UCU GGG GCA UGU UGG GUG GGC GUG CAA GGG GGU CCU GGG ACU CCU GAG UUU CUC AGG CC-3’, “UG high” 5’-6-FAM-UGC UGG GAU GGU CAU GGU GGG UGA CCC UGG GAU GGC CGC GGU GGG UGA CCC UGG AAU GGU-3’, “UG low” 5’-6-FAM-ACA GUC CAG UCG CAG CAG ACU CCC UCA CAG CAC CCA CCU CUC CCA CAA UCG GAU CCC AGG-3’. 5’ 6-FAM labelled double stranded DNAs were purchased from IDT corresponding to the UG high and UG low RNA sequences. Briefly, MYCN was titrated and mixed with 5 nM of fluorescently labelled RNA in a final buffer containing 50 mM NaCl, 20 mM Tris-HCl pH 7.9, 5% (v/v) glycerol, 1 mM DTT. Reactions were incubated at room temperature in the dark for 20 min. 18 µL of the reaction was transferred to a 384-well plate (Corning) and fluorescence anisotropy was detected using a Tecan Spark plate reader with a gain of 150, excitation of 470 ± 20 nm and emission of 518 ± 20 nm, Z-height of 15000 µm. Fluorescence anisotropy values were normalized by subtracting the anisotropy detected for the RNA alone from each well with protein and RNA. Experiments were performed at least 3 times. Data were graphed in GraphPad Prism (version 9.5.1) and fit using a single-site quadratic binding equation.

### Peptide Library Synthesis and Microarray Printing

MYCN (MYCN_HUMAN, Uniprot P04198) residues 1-160 were displayed as an overlapping peptide library of 72 20mers with an overlap of 18 aa and offset of 2 aa. µSPOT peptide array synthesis was carried out as previously described^59^. Peptides were synthesized on custom-prepared cellulose and laser cut towards cellulose discs with 4mm diameter pretreated to contain 9-fluorenylmethyloxycarbonyl-β-alanine (Fmoc-β-Ala) linkers at average loading density of 130nmol per disc. Synthesis was conducted using a MultiPep RSi robot (CEM GmbH). All chemicals were purchased at peptide synthesis purity or highest available purity (IRIS Biotech GmbH) with freshly prepared reagents. Fmoc deprotection was done using 20% piperidine in dimethylformamide (DMF), followed by washing with DMF and ethanol (EtOH). Peptide elongation was carried out by preactivated Fmoc protected AAs (0.5 M), coupling agent N,N′-diisopropylcarbodiimide (1 M) and ethyl 2-cyano-2-(hydroxyimino)acetate (1 M). Each coupling was performed three-times for 30 min, followed by capping with 1:25 acetic anhydride in DMF and washes with DMF and EtOH. After final deprotection with 20% piperidine, washed and dried discs were transferred to a 96-well plate. AA sidechain protecting groups were removed using 90% trifluoracetic acid (TFA), 3% triisopropylsilane (TIPS), 2% dichloromethane (DCM) and 5% water for 2 h while shaking at 1000 rpm. After removing the deprotection mix, peptides were left overnight to dissolve in a mixture containing 88.5% TFA, 4% trifluoromethanesulfonic acid, 5% water and 2.5% TIPS (200 µL/well) while shaking at 500 rpm. Resulting peptide cellulose conjugates (PCCs) were precipitated with 0.7 mL cooled diethyl ether, centrifuged for 10 min at 2000 g at 4 °C, followed by two additional diethyl ether washes. After drying pellets were dissolved in 250 μL DMSO per well. PCCs were transferred to 384 well plates and diluted 1:2 with filtered saline-sodium citrate buffer (150 mM NaCl, 15 mM trisodium citrate, pH 7.0) and printed as duplicates of 50 nl drops on coated blank slides (76 mm x 26 mm, Intavis Peptide Services GmbH) as microarrays using a SlideSpotter (CEM GmbH) followed by drying overnight.

### Microarray Binding Assay

Microarrays were equilibrated using PBST (137 mM NaCl, 2.7 mM KCl, 10 mM Na_2_HPO_4_, 1.8 mM KH_2_PO_4_, pH 7.4, 0.1 % v/v Tween20) for 30 min with slow rotation. After draining equillibration solution the Cy5-RNA-probe was incubated on the microarrays for 15 min at a concentration of 4 nM in PBST. Slides were subsequently washed three times with PBST and directly imaged on an Amersham Image Quant 800 (Cytiva) via Cy5 channel fluorescence detection with an exposure time of 5 sec. Throughout the assay, slides were protected from light using aluminum foil. Binding intensities were quantified using FIJI with the microarray profile addon (OptiNav). For each peptide/RNA binding event duplicate signals from 3 arrays, thus six raw grey intensities were obtained. Binding intensities for each array duplicate are averaged. Derivations were determined by measuring three independently processed and analyzed arrays.

### Total and nuclear protein isolation

For total native protein isolation, a HEPES lysis buffer (20 mM HEPES pH 7.9, 150 mM NaCl, 2 mM MgCl_2_, 0.2% v/v NP-40, 0.5 mM EDTA, 10% v/v glycerol) was used. Lysates were sonicated four times using 5 s pulses at 20% output with 10 s pauses in-between. Subsequently, 50 U/ml benzonase (Merck) or DNase I (Invitrogen) with murine RNase inhibitor (NEB) were added and the lysates were incubated under rotation at 4°C for 1 h before being cleared by centrifugation. For total denatured protein isolation, cells were lysed in RIPA lysis buffer (50 mM HEPES pH 7.9, 140 mM NaCl, 1 mM EDTA, 0.1% v/v SDS, 1% v/v Triton X-100, 0.1% w/v sodium deoxycholate). The lysates were incubated for 30 min at 4°C under rotation and subsequently cleared by centrifugation.

For nuclear protein isolation, cells were lysed in PIPES lysis buffer (5 mM PIPES pH 8.0, 85 mM KCl, 0.5% v/v NP-40). Lysates were then incubated on ice for 10 min, followed by centrifugation at 500 g for 10 min at 4°C to collect the nuclei. The supernatant containing the cytoplasmic proteins was discarded, and the nuclei were resuspended in HEPES lysis buffer to follow the procedure for total native protein isolation.

Phosphatase and protease inhibitors (1:1000; Sigma-Aldrich) were added to all lysis buffers. Protein concentrations were determined with the BCA or Bradford assay.

### Co-immunoprecipitation and immunoblotting

For co-immunoprecipitation (coIP), a total volume of 20 µl 1:1 A/G Dynabeads (Thermo Fisher Scientific) was used per IP. The beads were washed three times with 5 mg/ml BSA/PBS and then incubated overnight at 4°C under rotation in 5 mg/ml BSA/PBS containing 2.5 µg antibody against MYCN (Santa Cruz Biotechnology), HA-Tag (Merck), ZCCHC8 (Thermo Fisher Scientific) or PABPN1 (Proteintech). The beads were then washed three times with BSA/PBS and incubated for 6 h at 4°C with 2-4 mg of protein lysate under rotation. For glycerol gradient coIPs, samples were precleared by first incubating them with A/G beads alone for 1 h at 4°C, then the supernatant was added to the antibody-bound beads. After four washes in HEPES lysis buffer, samples were eluted in 2x Laemmli buffer and heated at 95°C for 5 min. Subsequently, the supernatants were loaded onto Bis-Tris gels and transferred to PVDF membranes (Merck) via wet transfer. Following blocking with 5% w/v skimmed milk or 5% v/v BSA in TBS-T, membranes were incubated with the appropriate primary antibodies overnight at 4°C. The membranes were probed with HRP– or IRDye-conjugated secondary antibodies that matched the host organism of the primary antibodies for 1 h at room temperature (RT). The LAS4000 Mini Imaging system (Fuji), Fusion FX (VILBER) or the Odyssey CLx (LI-COR Biosiences) were used to acquire the images.

### Glycerol gradient fractionation and precipitation

Nuclear protein lysates were loaded on a linear 10-30% glycerol gradient in 1x gradient buffer (20mM HEPES/KOH pH 7.9, 150 mM NaCl, 2 mM MgCl_2_, 0.2% v/v NP-40, 0.5 mM EDTA) and separated by ultracentrifugation in a Beckman Optima L-90 LK ultracentrifuge (Beckman Coulter) for 16 h at 41,000 rpm and 4°C using a SW41 rotor (Beckman Coulter). Manual fractionation of the gradients was performed by removing 500 ml off the top of the gradient and the fractions were either subjected to co-immunoprecipitation or precipitated with sodium deoxycholate (DOC) and trichloroacetic acid (TCA). For DOC/TCA precipitation, 0.15% w/v DOC in H_2_O was added to each glycerol fraction and the samples were incubated for 10 min at RT. After centrifugation at 12,000 rpm for 15 min at 4°C, the resulting pellet was washed with ice-cold acetone and centrifuged at 12,000 rpm for 10 min at 4°C. The pellets were then left to dry and resuspended in 2x Laemmli buffer for immunoblotting.

### MYCN glycerol gradient LFQ interactome in asynchronous and S phase cells

Cells were synchronized at the S phase as described above. S phase and asynchronous nuclear lysates were loaded on 10-30% glycerol gradients and MYCN immunoprecipitation (IP) was performed using pooled fractions 12-19 for both conditions. IgG IPs were used as controls. Bound proteins were eluted in 2x NuPAGE LDS Sample Buffer (Life Technologies) supplemented with 1 mM DTT, heated at 70 °C for 10 min, alkylated with 5.5 mM chloroacetamide for 30 min in the dark, and separated by SDS-PAGE on a 4-12% gradient Bis–Tris gel. Proteins were fixed with 50% methanol, 10% acetic acid and stained using the Colloidal Blue Staining Kit (Life Technologies). The gel was cut into 4 similarly sized fractions, which were cut into 1 × 1 mm cubes, destained in 50% ethanol, 25 mM ammonium bicarbonate and digested in-gel using 0.625 µg of MS-approved trypsin (Serva) per fraction overnight at 37°C^60^. Peptides were extracted from the gel using a series of increasing acetonitrile percentages and desalted using reversed-phase C18 StageTips^61^.

### MYCN glycerol gradient TMT interactome after EXOSC10 depletion

Control and EXOSC10 depleted nuclear lysates were loaded on 10-30% glycerol gradients and MYCN immunoprecipitation (IP) was performed using pooled fractions 13-20 for both conditions. Dried beads were resuspended in 2x NuPAGE LDS Sample Buffer (Life Technologies) supplemented with 2 mM DTT and boiled for 15 min at 70°C, then alkylated with 10 mM chloroacetamide for 30 min in the dark at room temperature. To remove detergents, proteins were purified on SP3 beads^62^. Briefly, hydrophilic and hydrophobic Sera-Mag magnetic carboxylate modified particles (Sigma) were combined 1:1 (V:V), washed in water and resuspended in water at 20 mg/ml. 20 µl of combined beads were added to each sample. Absolute ethanol was added to 50% (V:V) and samples were shaken for 5 min at 1000 rpm. After magnet binding, supernatants were removed and the beads were washed three times with 80% ethanol. Proteins were digested on-bead in the digestion buffer (150 mM HEPES, pH 8) with 0.5 µg of MS-approved trypsin (Serva) per sample overnight at 37°C with shaking. After magnet binding, the supernatants were collected, and the beads were washed twice with the digestion buffer. Wash supernatants were combined with respective peptide supernatants. Digestion was stopped by adding formic acid (FA) to 1%, and precipitates were removed by centrifugation after a 30 min incubation at 4°C. Supernatants were desalted using reversed-phase C18 StageTips2 and eluted in 50% acetonitrile (ACN), 0.1% FA. Eluates were vacuum-centrifuged until completely dry and reconstituted in 30 µl of 50 mM HEPES pH 8.5, 30% ACN. Samples were barcoded with 0.1 µg of ACN-dissolved TMTPro labels (Thermo Scientific) for 1 h and quenched with 5% hydroxylamine for 15 min^63^. An equal aliquot (5% vol) of each sample was mixed and the ratio check was performed as described^64^. Samples were vacuum-centrifuged until completely dry and stored at –80°C. After ratio check, samples were reconstituted in 30 µl of 1% trifluoroacetic acid and masses adjusted treatment-wise before mixing. The mixed sample was fractionated into 8 fractions ranging from 2% to 50% ACN using the Pierce High pH Reversed-Phase Peptide Fractionation Kit (Thermo Scientific), according to manufacturer’s instructions. Column-eluted samples were completely vacuum-dried and resuspended in 80% ACN, 0.1% FA, then vacuum-dried to 5 µl to remove ACN, and volumes were adjusted with 1-2 µl of 0.1% FA for MS loading.

### Quantitative mass spectrometry

Samples were analyzed on a quadrupole Orbitrap mass spectrometer (Exploris 480, Thermo Scientific) equipped with a UHPLC system (EASY-nLC 1200, Thermo Scientific). They were loaded onto a C18 reversed-phase column (55 cm length, 75 mm inner diameter, packed in-house with ReproSil-Pur 120 C18-AQ 1.9-mm beads, Dr. Maisch GmbH) and eluted with a gradient from 2.4 to 32% ACN (LFQ run) or 2.4 to 33.6% ACN (TMT run) containing 0.1% FA in 90 min (LFQ run) or 120 min (TMT run). The mass spectrometer was operated in data-dependent mode, automatically switching between MS and MS2 acquisition. Survey full scan MS spectra (m/z 300–1,650, resolution: 60,000, target value: 3 × 106, maximum injection time: 28 ms) were acquired in the Orbitrap. The 15 (LFQ run) or 20 (TMT run) most intense precursor ions were sequentially isolated, fragmented by higher energy C-trap dissociation (HCD) and scanned in the Orbitrap mass analyzer (normalized collision energy: 30% (LFQ run) or 33% (TMT run), resolution: 15,000, target value: 1 × 105, maximum injection time: 40 ms, isolation window: 1.4 m/z (LFQ run) or 0.8 m/z (TMT run)). Precursor ions with unassigned charge states, as well as with charge states of +1 or higher than +6, were excluded from fragmentation. Precursor ions already selected for fragmentation were dynamically excluded for 25 s (LFQ run) or 30 s (TMT run). For the TMT run, TurboTMT scan feature (TMTPro reagent) was implemented.

### Proximity ligation assay (PLA)

Cells were seeded in 384-well plates (Perkin Elmer). Following treatment, cells were fixed with 4% methanol-free paraformaldehyde (Electron microscopy science) for 20 min at RT, permeabilized with 0.1% v/v Triton-X for 10 min at RT, and blocked with 5% w/v BSA/PBS for 30 min at RT. Cells were incubated with the appropriate mouse and rabbit antibody pairs in blocking buffer overnight at 4°C. PLA was performed using the Duolink In Situ Kit (Sigma-Aldrich) according to the manufacturer’s instructions. Briefly, after washing with TBS-T, cells were treated with plus and minus PLA probes specific for rabbit and mouse antibodies and incubated at 37°C for 1 h. After washing with PLA wash buffer A (Sigma-Aldrich), ligation was performed at 37°C for 30 min. Following washing with PLA Wash Buffer A, *in situ* PCR amplification was performed by incubating cells with Alexa 568 or 644-conjugated oligonucleotides for 90 min at 37°C in the dark. The cells were washed with PLA wash buffer B (Sigma-Aldrich) and the nuclei counterstained with Hoechst 33452 (Sigma-Aldrich) for 10 min at RT in the dark. Subsequently, the cells were washed and overlaid with PBS. 30 to 60 image fields per well were acquired at 40x magnification, 100% excitation and non-confocal mode using the Operetta High Content Imaging System with a total of at least 250 cells per sample. Images were processed with the Harmony High Content Imaging and Analysis Software. Conditions were compared using the GraphPad Prism (v10.0.0) software and *p*-values were calculated with the Mann-Whitney test.

### Electrophoretic mobility Shift Assay (EMSA)

His-MBP-tagged MYCN Full-Length (1-464) was incubated with 5 nM 6-FAM-labeled UG-High RNA purchased from IDT (sequence 5’-6-FAM-UGC UGG GAU GGU CAU GGU GGG UGA CCC UGG GAU GGC CGC GGU GGG UGA CCC UGG AAU GGU) for 20 min at room temperature in a final buffer containing 50 mM NaCl, 18 mM HEPES pH 7.4, 5% glycerol, 1 mM DTT, 5 mM Tris-Cl pH 7.9. 2 μL of 80% glycerol was added prior to loading 10 μL of the binding mixture onto Native PAGE. The complexes were separated at 100 V on 3-12% Bis-Tris Native PAGE (Invitrogen) at 4°C for 90 min. FAM fluorescence was detected using a Typhoon FLA9500 imager (GE) (PMT 900). Each EMSA was performed in three independent replicates. To determine Kd based on Full-Length His-MBP-tagged MYCN EMSAs, the percent of RNA probe bound by the major RNA binding event indicated in Figure 3C was quantified using ImageQuant 7.0. The percent of probe bound was plotted using Prism version 9.5.1 and the Kd was calculated using a single-site quadratic binding equation.

### 4sU-RNA labeling and sequencing

Cells were treated as described in the main text. After 10 min incubation with 1 mM 4-thiouridine (4sU, Sigma-Aldrich), cells were incubated for 2 h in non-4sU-containing medium (chase). Cell number was counted in all conditions and lysis was performed using QIAzol reagent (QIAGEN). Cells were spiked based on their cell number with 4sU-labeled murine NIH3T3 cells. Total RNA was extracted using the miRNeasy kit (QIAGEN), including an on-column DNase digestion step and RNA quantity and quality were assessed using a Nanodrop spectrophotometer. 4sU RNA enrichment was performed by mixing 60 µg total RNA with EZ-Link Biotin-HPDP (Pierce) in 0.2 mg/ml dimethylformamide and biotin labelling buffer (10 mM Tris pH 7.4, 1 mM EDTA), followed by incubation for 2 h at RT under rotation. Biotinylated RNA was then purified with chloroform-isoamyalcohol extraction using MaXtract high density tubes (QIAGEN). The upper aqueous phase was collected and 1/10 volume of 5 M NaCl and an equal volume of isopropanol were added. Samples were centrifuged at 20,000g for 20 min at 4°C. The RNA pellets were washed with 75% v/v ethanol and centrifuged at 20,000g for 10 min at 4°C before being left to dry and resuspended in nuclease-free water. Biotinylated RNA was pulled down by mixing the RNA with Dynabeads MyOne Streptavidin T1 beads (Invitrogen) in Dynabeads wash buffer (2 M NaCl, 10 mM Tris pH 7.5, 1 mM EDTA, 0.1% v/v Tween 20) for 15 min at RT under rotation. After washing, 4sU-labeled RNA was eluted from the beads with 100 µl freshly prepared 100 mM DTT in nuclease-free water and purified using the RNeasy MinElute Kit (QIAGEN). Samples were quantified using the RiboGreen Assay (Invitrogen) and 60 ng 4sU RNA per sample were used for cDNA library preparation. The NEBNext rRNA Depletion Kit Human/Mouse/Rat (New England Biolabs; NEB) was first used to degrade rRNA according to manufacturer’s instructions. Libraries were then generated using the NEBNext Ultra II Directional RNA Library Prep Kit (NEB) following manufacturer’s instructions. The optimal number of PCR cycles for cDNA library amplification was determined via qPCR. A 1/10 dilution of each sample was mixed with PowerUP SYBR Green Master Mix (Thermo Fisher Scientific) and Illumina compatible primers in 10µl reactions. Reactions were run at a StepOnePlus Real-Time PCR System (Thermo Fisher Scientific). Optimal PCR cycles for each sample were determined by subtracting the resulting Ct values by 4. All samples were then amplified for 10 PCR cycles and size selected using SPRIselect beads (Beckman Coulter).

### Enhanced cross-linking and immunoprecipitation (eCLIP)

eCLIP was performed as described in ^31,65^. All described eCLIPs were performed in biological duplicate with one corresponding pooled size-matched input. 30-40 million cells at 70% confluence per replicate were treated as described in main text. Cells were then crosslinked with UV light at 254 nm for 2 min at 400 mJ/cm^2^. Non-UV crosslinked cells were used as negative controls. All cells were lysed in cold eCLIP lysis buffer (50 mM Tris-HCl pH 7.4, 100 mM NaCl, 1% v/v Igepal CA-630, 0.1% v/v SDS and 0.5% w/v sodium deoxycholate) supplemented with protease and phosphatase inhibitors and murine RNase inhibitor (New England Biolabs). Lysates were sonicated with a M220 Focused-ultrasonicator (Covaris) for 3 min at low settings and then treated with Turbo DNase and RNase I for 5 min at 37°C. Following centrifugation at 15,000g for 10 min at 4°C, the supernatants were added to A/G Dynabeads (Thermo Fischer Scientific) pre-coupled with 15 µg MYCN antibody (Santa Cruz Biotechnology), HA antibody (Merck) for the EXOSC10 mutant or murine IgG Control antibody (Sigma-Aldrich). Samples were incubated for 3 h at 4°C under rotation. Afterwards, 2% v/v of the lysates, including beads, were kept as sized-matched input samples. All samples were washed using wash buffer (20 mM Tris-HCl pH 7.4, 10 mM MgCl2, 0.2% v/v Tween-20 and 5 mM NaCl) and high-salt wash buffer (50 mM Tris-HCl pH 7.4, 1 M NaCl, 1% v/v Igepal CA-630, 0.1% v/v SDS and 0.5% w/v sodium deoxycholate). RNA was dephosphorylated with FastAP thermosensitive alkaline phosphatase (Thermo Fisher Scientific) and T4 polynucleotide kinase (New England Biolabs). On-bead 3’ RNA adapter ligation was carried out for the IP samples using T4 RNA ligase 1 (New England Biolabs) and X1A/X1B or X2A/X2B RNA adapter pairs (sequences and modifications in KRT). IP and input samples were then mixed with 4x NuPAGE LDS buffer and DTT, loaded run on 8% Bis-Tris gels and transferred to nitrocellulose membranes (Merck) overnight at 4°C at 30 V. Membranes were cut out at the observed protein size and extending to 75 kDa above that point. All membrane slices were then incubated with proteinase K (New England Biolabs) at 37°C for 20 min followed by a 20 min incubation at 50°C. RNA was purified using the RNA Clean and Concentrator-5 kit (Zymo Research), input samples were dephosphorylated and 3’RNA Ril19 adapters were appended. Samples were reverse transcribed using an AR17 primer and the AffinityScript enzyme (Agilent). Remaining primer and dNTPs were removed with ExoSAP-IT (Thermo Fisher Scientific) and RNA was degraded by incubating with 100 mM NaOH at 70°C for 10 min. Samples were purified with MyOne Silane Dynabeads (Thermo Fisher Scientific) before 5’ ligation of the 10N randomer-containing adapter (Rand3Tr3). The cDNA samples were purified using MyOne Silane Dynabeads and optimal PCR cycle number was determined as for 4sU-seq. All eCLIP required <16 cycles to amplify, with only the IgG eCLIP libraries requiring 20 cycles. Final PCR amplification was carried out using the Q5 High-Fidelity 2x PCR Master Mix (New England Biolabs). Libraries were purified using SPRIselect beads (Beckman Coulter) and size selected by loading onto a 3% agarose gel. Extracted gel slices at 175-350 bp were purified using the QIAquick Gel Extraction Kit (QIAGEN).

### Spike-in chromatin immunoprecipitation and sequencing (ChIP-Rx)

Following treatment, formaldehyde was added to the cell culture medium to a final concentration of 1% v/v, and incubated for 5 min rotating at RT to crosslink protein to DNA. The reaction was stopped by adding glycine to a concentration of 125 mM, and incubating for 5 min under constant rotation. Subsequently, cells were washed and harvested in ice-cold PBS containing protease and phosphatase inhibitors (Sigma Aldrich), which were added to all buffers mentioned hereafter. Cells were pelleted (1,500 rpm, 20 min, 4°C) and resuspended in Lysis buffer I (5 mM PIPES pH 8.0, 85 mM KCl, 0.5% NP-40). 10 % of murine spike-in cells were added based on cell number. After 20 min incubation on ice, nuclei were collected by centrifugation at 1,500 rpm for 20 min at 4°C. Pelleted nuclei were lysed by incubation in Lysis buffer II (10 mM Tris pH 7.5, 150 mM NaCl, 1 mM EDTA, 1% NP-40, 1% sodium deoxycholate, 0.1% SDS) for 10 min on ice. The released chromatin was fragmented by using the Covaris Focused Ultrasonicator M220 (50 min; cycles/burst, 200; peak power, 75; duty factor, 10). The correct fragment size of 150-300 bp was validated by agarose gel electrophoresis. The fragmented chromatin was centrifuged at 14,000 rpm for 20 min at 4°C. Per IP, 100 µl of each Protein A and Protein G-coupled Dynabeads (Thermo Fisher Scientific) were pre-incubated with 15 µg MYCN antibody (sc-56729, Santa Cruz Biotechnology) in the presence of 5 mg/ml BSA overnight at 4°C. Protein-coupled beads were added to the chromatin and incubated for at least 6 h at 4°C on a rotating wheel. Immunoprecipitated protein-chromatin complexes were washed 3 times each with ice-cold Wash buffer I (20 mM Tris pH 8.1, 150 mM NaCl, 2 mM EDTA, 1% Triton X-100, 0.1% SDS), Wash buffer II (20 mM Tris pH 8.1, 500 mM NaCl, 2 mM EDTA, 1% Triton X-100, 0.1% SDS), Wash buffer III (10 mM Tris pH 8.1, 250 mM LiCl, 1 mM EDTA, 1% NP-40, 1% sodium deoxycholate; including a 5 min incubation at 4°C), and once with ice-cold TE buffer. Chromatin was eluted twice by incubating in 150 µl Elution buffer (100 mM NaHCO3, 1% SDS) for 15 min at RT. Eluted IP samples and respective input samples were first depleted of RNA by incubation with RNase A for 1 h at 37°C, de-crosslinked overnight at 65°C shaking, and finally digested with Proteinase K for 2 h at 45°C with shaking. DNA was isolated by phenol-chloroform extraction followed by ethanol precipitation. The amount of DNA was quantified using Quant-iT PicoGreen dsDNA assay (Thermo Fisher Scientific) and libraries were generated using the NEBNext Ultra II DNA Library Prep Kit for Illumina (New England Biolabs) following manufacturer’s instructions.

### Cleavage Under Targets and Release Using Nuclease (CUT&RUN)

CUT&RUN followed by sequencing was performed as previously described in^66^. In brief, adherent cells were harvested using Accutase (Sigma) and 10% murine NIH 3T3 cells were added as spike-in. Cells were washed by adding 1 ml of Wash buffer (20 mM HEPES pH 7.5, 150 mM NaCl, 0.5 mM Spermidine) followed by centrifugation (3 min, 600 g, RT). After three repetitions, cells were coupled to Concanavalin A-coated magnetic beads (Polysciences Europe) by incubating in 1 ml Wash buffer supplemented with 10 µl bead slurry per sample (pre-washed with 1 ml Binding buffer (20 mM HEPES pH 7.5, 10 mM KCl, 1 mM CaCl_2_, 1 mM MnCl_2_)) for 10 min rotating at room temperature (RT). After magnetically separating the bead-coupled cells, the supernatant was removed and replaced by 150 µl Dig-wash buffer (Wash buffer supplemented with 0.05% Digitonin) containing 2 mM EDTA to permeabilize the cells. 3 µg of each, MAX (Proteintech), MNT (Thermo Fisher Scientific), or SIN3A (Novus Biologicals) antibody were added per sample and incubated with shaking (800 rpm) at 4°C overnight. The next day, cells were washed two times with 1 ml Dig-Wash buffer, followed by incubation with Protein A/G-MNase fusion protein at a concentration of 1 µg/ml diluted in 150 µl Dig-Wash buffer for 1 h shaking (800 rpm) at 4°C. Cells were washed two times with 1 ml Dig-Wash buffer followed by a wash with 1 ml Low salt rinse buffer (20 mM HEPES pH 7.5, 0.05% Digitonin, 0.5 mM Spermidine). After removing the Low salt rinse buffer, the MNase coupled to the primary antibodies was activated by adding 200 µl of ice-cold Incubation buffer (3.5 mM HEPES pH 7.5, 10 mM CaCl_2_, 0.05% Digitonin) and incubated for 30 min on ice. To stop the reaction, 200 µl of STOP buffer (170 mM NaCl, 20 mM EGTA, 0.05% Digitonin, 50 µg/ml RNAse A, 25 µg/ml Glycogen) were added and samples were incubated for 30 min at 37°C to release the MNase-cleaved DNA fragments from the insoluble nuclear chromatin. To digest the protein, SDS and proteinase K were added to a concentration of 0.1% and 250 µg/ml, respectively, and incubated at 50°C for 1 h. DNA was isolated by phenol-chloroform extraction and quantified by determining the concentration of small fragments (7-50 bp) using the Fragment Analyzer (Agilent). The amount of input material was equalized, and libraries were prepared using the NEBNext Ultra II DNA Library Prep Kit for Illumina (New England Biolabs).

### mRNA sequencing

After treatment, cells were washed with PBS and lysed with RLT buffer (QIAGEN) supplemented with beta-mercaptoethanol. Total RNA was extracted with RNeasy mini columns (QIAGEN) including an on-column DNase I digestion step according to manufacturer’s instructions. RNA concentration was determined using a Nanodrop and RQN values were determined with a Fragment Analyzer (Agilent). RNA samples with RQN > 9 were used for mRNA isolation with the NEBNext Poly(A) mRNA Magnetic Isolation Module (New England Biolabs). cDNA libraries were then prepared using the NEBNext Ultra II Directional RNA Library Prep Kit for Illumina (New England Biolabs) according to manufacturer’s instructions. Optimal PCR cycle number was assessed via qPCR as with 4sU and eCLIPseq, Libraries were amplified for 10 PCR cycles and size selected using SPRIselected beads (Beckman Coulter).

### 3’RNA sequencing

3’RNAseq libraries were generated as described in ^67^. Cells were incubated with 1 mM 4sU for 10 min, then immediately lysed and spiked with 4sU-labelled murine NIH3T3 cells based on cell number. Total RNA and 4sU RNA isolation was performed as for 4sU-seq. 4sU RNA (200 ng) was then *in vitro* polyadenylated in 20 µl reactions using a poly(A)-tailing kit (Thermo Fisher Scientific) in the presence of murine RNase inhibitor (NEB). A lower amount of *E.coli* poly(A) polymerase (0.4 U) was used and the reactions took place at 30°C for 30 min to generate short polyA tails. Samples were then purified using the RNA Clean and Concentrator-5 kit (Zymo Research). 200 ng of samples with (all RNAs) and without (only polyA) *in vitro* polyadenylation were then rRNA depleted using the NEBNext rRNA Depletion Kit Human/Mouse/Rat (New England Biolabs) according to manufacturer’s instructions. Libraries were generated using Quant Seq 3’mRNA-Seq REV library prep kit (Lexogen) following manufacturer’s instructions. Optimal PCR cycles were determined as described in 4sU-seq. All samples were amplified for 16 cycles and bead purified.

### High-throughput sequencing

Libraries were sequenced for 60 cycles per read in case of paired-end sequencing and for 100 cycles in case of single-read sequencing on a NextSeq2000 System (Illumina) according to the manufacturer’s instructions. All samples were paired-end sequenced, except for the 3’RNAseq samples, which were single-read sequenced with a Custom Sequencing Primer (Lexogen).

### Quantification and statistical analysis FASTQ file generation

For all sequenced libraries, base calling was performed with Illumina’s BCL Convert Software, implemented in the Illumina Dragen software (v3.7.4 or v3.10.4) and high-quality PF-clusters (according to the chasity filter) were selected for further analyses. Overall sequencing quality was determined with FastQC (v0.11.9).

### ChIP-Rx and CUT&RUN data processing

Human reads were mapped to hg19 and mouse reads were mapped to mm39 using Bowtie2 (v2.3.5.1) with the –N 1 option. For MYCN ChIP-Rx and MAX CUT&RUN BAM files, normalization ratios were calculated by dividing the mm39 reads in each sample by the lowest number of mm39 reads. The normalization ratios were then used to multiply the corresponding hg19 reads and derive nomalized read numbers. MNT and SIN3A CUT&RUN BAM files were normalized to sequencing depth. Normalized BAM files were sorted and indexed using Samtools (v1.7) and converted to bigwig using deeptools (v3.5.1) bamCoverage. Bigwig files were loaded on Integrated Genome Browser (9.1.10) to inspect read distribution. Matrices of counts around the transcription start sites of IMR-32 expressed genes were generated with deeptools computeMatrix. The resulting matrices were opened in R (4.2.2) to calculate the 98^th^ percentile of values. Furthermore, *p*-values for each bin were calculated in R using two-sided Wilcoxon tests and *p*-value heatmaps were then generated using pheatmap (1.0.12). To perform 2% trimming, all values above the calculated 98^th^ percentiles were filtered using deeptools computeMatrixOperations filterValues and the matrices were then loaded on deeptools plotProfile to generate density plots. Log_2_ fold-changes (FC) were calculated usind deeptools bigwigCompare adding a pseudocount of 1 and matrices were again generated with computeMatrix. MYCN ChIP-Rx peaks were called using MACS3 (v3.0.0a7) with a *p*-value < 0.001 threshold.

### mRNAseq data processing

mRNA reads were aligned to hg19 using the STAR aligner (v2.7.10a) and a splice junction-aware index ^68^. BAM files were sorted with Samtools and loaded on featureCounts (v2.0.3) with options –p –O –-minOverlap 10 –B –C to count reads falling in hg19 genes. DESeq2 (v1.38.3) was used to calculate log_2_ fold-changes and *q*-values between the conditions and ggplot2 (v3.4.2) was used to generate volcano plots based on the DESeq2 output. A threshold of at least 10 reads in all samples was set to generate a list of IMR-32 expressed genes. Transcript per million (TPM) values were derived by dividing the mean of all control sample reads in each gene by the corresponding length (RPK; Reads per Kilobase) and then dividing each individual RPK by the RPK sum and multiplying by a million. The gene list containing TPM values was sorted in ascending order and split into 30 bins of ca. 505 genes with increasing TPM. The ChIP-Rx and CUT&RUN log_2_FC matrices from deeptools were opened in R, the values ± 750 bp around the TSS were kept and sorted according to increasing TPM. The bins were then plotted using ggplot2 and DescTools (0.99.49) to show confidence intervals for each bin and condition.

The size factors from DESeq2 were first applied to the RNAseq reads before loading them on GSEA (v4.3.2) for gene set enrichment analysis with 1000 gene set permutations.

### 4sU-RNAseq data processing

Human reads were mapped to hg19 and mouse reads were mapped to mm39 using the STAR aligner (v2.7.10a) and a splice junction-aware index. Exonic, rRNA and blacklisted^69^ regions intersecting the generated BAM files were removed using bedtools. Spike-in normalization was performed in the same way as for ChIP-Rx and CUT&RUN. BAM files were sorted with Samtools and loaded on featureCounts (v2.0.3) with options –-minOverlap 10 –B –C to count reads falling in non-coding RNA regions (PROMPT, eRNA, lncRNA, snoRNA hosting intron, and regular intron). PROMPT annotations were taken from ^67^ and lifted over to hg19 using the UCSC liftOver tool. lncRNA, snoRNA and intron coordinates were downloaded from UCSC table browser based on Ensembl annotations. Only the longest lncRNAs were kept when multiple lncRNAs shared the same start or end coordinate and all lncRNAs overlapping more than 50% with annotated introns were removed from further analysis using bedtools (v2.26.0) intersect. snoRNA hosting introns were derived by intersecting snoRNA with intron coordinates using bedtools. All introns below 10 bases were removed and only the longest introns per gene were kept. eRNA annotations were taken from ^70^. The featureCounts output files were loaded on DESeq2, normalization factors were set to 1 for all samples and volcano plots were generated with ggplot2 using the DESeq2 outputs. Box plots were generated based only on non-coding RNAs that had *p*-value < 0.05 and log_2_FC > 0.5 or log_2_FC < –0.5.

### eCLIP data processing

eCLIP samples were processed based on paired-end eCLIP analysis protocols ^31,65^. Adapters were trimmed twice using cutadapt (v1.14). Trimmed FASTQ files were then first aligned to a genome index consisting only of Repbase annotated repetitive elements using the STAR aligner (v2.7.6a). Reads that did not map to repetitive elements were then aligned to hg19 using STAR. Genome mapped BAM files were sorted with Samtools and PCR duplicates were removed by a custom python script ^31^. BAM file read pairs were then sorted, indexed and merged using Samtools and then converted to sequence depth normalized, stranded bigwig files using a custom python script ^31^. The CLIPper ^31,65^ (v5d865bb) peak calling algorithm was used to determine high-confidence peaks in the BAM files. The thresholds used were log_2_FC > 3 reads over the size-matched input and *p*-value < 0.001. A final list of high-confidence eCLIP peaks merging both biological duplicates was generated by first normalizing all BAM files to sequencing depth using Samtools, removing overlapping regions with a custom perl script ^31,65^ and then running IDR (irreproducible discovery rate) (v2.0.2) on both replicates with a 0.01 cutoff. The calculated log_2_FC over input was normalized based on the new IDR positions and the final list of eCLIP peaks was generated using custom perl scripts ^31,65^.

All eCLIP peak lists were filtered based on the ENCODE blacklist ^69^ and a manually curated eCLIP blacklist (ENCODE, ENCFF039QTN) using bedtools. The IgG eCLIP and U2OS MYCN eCLIP lists were used as additional blacklists and used to further filter the eCLIP peaks from MYCN and EXOSC10 mutant eCLIPs. Finally, all MYCN eCLIPs found 500 bp upstream or downstream of called MYCN ChIP-Rx summits (see ChIP-Rx data processing) were also filtered using bedtools window. Bookended peaks were merged in all samples using bedtools merge. For peak annotation, eCLIP peak files were sequentially intersected using bedtools with annotated introns, coding exons, 5’ and 3’ UTR and TSS coordinates derived from the UCSC table browser and biomart hg19. Intronic peaks were removed when they would overlap with any other feature, the same procedure was then sequentially followed for 3’ UTR, 5’ UTR and coding exons. Any peaks not intersecting with these features were considered intergenic. Gene annotation of the eCLIP peaks was performed with HOMER^71^ (v4.11.1) using a GENCODE hg19 genome file. *De novo* motif enrichment analysis was also carried out with HOMER based on intronic MYCN eCLIP peaks and a custom background file containing intronic coordinates from IMR-32 expressed cells. Shuffled peak files were generated with bedtools shuffle using the –incl option to restrict the random sequences to IMR-32 expressed gene coordinates. Distances between closest peaks were calculated for each peak list using bedtools closest with –d –io –t first options. Adjacent duplicate values and all distances lower than 10 bp were removed. *P*-values between MYCN eCLIP peak lists and their shuffled counterparts were calculated with Wilcoxon rank-sum test in GraphPad Prism (v8.2.1). Nucleotide sequences of MYCN eCLIP peaks were derived with bedtools getfasta and the UG dinucleotide content was counted in Excel spreadsheets for both eCLIP and shuffled peak lists. *P*-values were calculated as for peak distance peak lists.

Sequence depth normalized BAM files were separated by strand using Samtools and converted to bigwig using deeptools. Bigwig files of merged duplicates on the same strand were made using deeptools bigwigCompare with the mean operation and stranded matrices were generated using deeptools computeMatrixOperations filterStrand. 2% trimming and density plot generation was carried out as described in ChIP-Rx. Heatmaps were generated using 2% trimmed matrices with deeptools plotHeatmap and sorted with the –-sortUsing mean and –-sortUsingSamples options.

### 3’RNAseq data processing

Single-read 3’RNAseq analysis was carried out as in ^72^. FASTQ files were trimmed with the bbduk.sh (v39.01) script from BBMAP software with the options k = 13 ktrim = r useshortkmers = t mink = 5 qtrim = t trimq = 10 minlength = 20. The adapter reference file contained Illumina TruSeq adaptor sequences 1-27 and an additional 18A sequence to trim homopolymeric A-tails. Trimmed reads were then mapped to hg19 and mm39 using the STAR aligner (v2.7.10a) and a splice junction-aware index. The resulting BAM files were sorted and indexed with Samtools and then spike-normalized as described for 4sU-seq, with the exception that the second lowest mouse reads were used for normalization. A custom python script with HTSeq (v0.13.5) was used to generate stranded bedgraph files that report the coverage of uniquely mapping 5’ ends (with the Lexogen REV kit, these correspond to RNA 3’ ends) ^72^. This process was carried out for both polyadenylated (all RNAs) and non-polyadenylated libraries (only polyA). Bedgraphs were converted to bigwig using the UCSC bedGraphToBigWig tool and biological duplicate bigwig files were merged as described before. Log_2_FC between bigwig files was calculated as before and density plots were generated with deeptools plotProfile.

### Mass spectrometry data analysis

Raw data files were analyzed using MaxQuant v1.5.2.8 (LFQ run) or v2.3.0.0 (TMT run) ^73,74^. Parent ion and MS2 spectra were searched against a reference proteome database containing human protein sequences obtained from UniProtKB version 2020_02 (LFQ run) or 2021_03 (TMT run) using Andromeda search engine^75^. Spectra were searched with a strict trypsin specificity and allowing up to two miscleavages. Cysteine carbamidomethylation was searched as a fixed modification, whereas protein N-terminal acetylation and methionine oxidation were searched as variable modifications. The dataset was filtered based on posterior error probability (PEP) to arrive at a false discovery rate of below 1% estimated using a target-decoy approach^76^. For the TMT run, TMTPro label correction factors were imported into MaxQuant according to manufacturer’s specifications. TMTPro reporters were searched at the MS2 reporter ion level with a mass tolerance 0.003 Da and with PIF ≥ 0.75.

Statistical analysis was performed using the R software environment (version 4.2.2). For the LFQ run, missing values were imputed using Perseus^77^ (version 1.6.10.0). Potential contaminants, reverse hits, hits only identified by site and hits with no unique peptides were excluded from the analysis. For the TMT run, corrected reporter intensities were normalized treatment-wise with the quantile normalization approach, using the Limma package^78^. Additionally, to account for different levels of immunoprecipitated MYCN and allow better assessment of differential binding, intensities were normalized per TMT channel by dividing them with the MYCN intensity from the respective channel. *P*-values and false discovery rates were calculated using a moderated t-test (Limma package)^78^. Volcano plots were generated in R using ggplot2 and enriched complexes were defined using the COMPLEAT online tool^79^.

### Western blot and cell viability quantification

To quantify MYCN protein levels in the glycerol gradient experiments, the scanned immunoblot images were loaded into ImageStudio Lite (v5.2.5; LI-COR Biosciences). For each gradient, the lowest signal was subtracted from all other values and then each value was divided by the highest signal to obtain relative MYCN intensities between 0-1. This was performed for at least three replicates of each gradient. *P*-values between the conditions in each fraction were calculated in GraphPad Prism (v8.2.1) using multiple unpaired t-tests.

The basic analyzer tool in the Incucyte live-cell Imaging software (v2021C) was used to quantify phase and fluorescent object metrics in real-time. The obtained cell confluence values (Mean ± SD) were visualized in growth curves as percentage over time using GraphPad Prism (v10.0.0). Measurement endpoint confluency (Mean ± SD) were visualized in a grouped barchart and *p*-values between different treatment conditions were calculated based on the endpoint values using paired Welch’s t-test.

## Supplemental Figure legends

**Figure S1: MYCN forms stable complexes in S phase with RNA-binding proteins. Related to Figure 1**.

**(A)** Propidium-Iodide (PI) stained FACS profiles of cell cycle-synchronized IMR-5 cells. Cells were synchronized using double thymidine block (2 mM thymidine) and released for 4 h (S phase), 8 h (G2/M phase) and 13 h (G1 phase) before staining with PI. Asynchronous cells are shown as a control (n=2).

**(B)** Calibration of the 10-30% glycerol gradient ultracentrifugation using bovine serum albumin (BSA), maltose binding protein (MBP), alcohol dehydrogenase (ADH) and apoferritin to approximate which fraction corresponds to which protein weight. Larger complexes sediment together in higher fractions and are only disassembled during the SDS gel electrophoresis (n=2).

**(C)** Immunoblot of endogenous MYCN, MAX, EXOSC10 and total RNAPII sedimentation patterns in 10–30% glycerol gradients from asynchronous or S phase synchronized IMR-5 cells. Nuclear lysates were treated with 50 U/ml benzonase in both conditions (n=4).

**(D)** Quantification of the normalized MYCN band intensities from the immunoblots in (D). *P*-values were calculated using multiple unpaired t-tests. Dots indicate mean and error bars depict range of values. (n=4; * indicates *p*-values < 0.05).

**(E)** MYCN protein interactions in IMR-5 cells comparing asynchronous and S phase synchronized cells. Nuclear lysates were loaded on 10-30% glycerol gradients and MYCN IP was performed using pooled fractions 12-19 for both conditions. IgG IPs were used as controls. The x-axis displays the enrichment (log_2_FC) of proteins in asynchronous cells compared to the IgG control, the y-axis displays the log_2_FC of proteins in S phase synchronized cells compared to the IgG control. A threshold of log_2_FC > 0.25 was set for enrichment. Proteins enriched in S phase are marked in light green, S phase enriched RNA-binding proteins (RBP) are highlighted in dark green (n=1).

**Figure S2: EXOSC10 depletion shifts MYCN interactions. Related to Figure 1**.

(**A**) Volcano plots showing log_2_ fold-change (FC) (x-axis) and the statistical significance (y-axis) of 4sU-labeled RNA levels between shEXOSC10 and control IMR-32 cells, treated as in Figure 1A. Each point represents an annotated enhancer RNA (eRNA), long non-coding RNA (lncRNA), snoRNA-hosting introns or regular introns (from left to right). Only RNAs with *p*-value < 0.05 and log_2_FC > 0.5 or < –0.5 were considered significantly altered (n=3).

(**B**) Quantification of nuclear proximity ligation assay (PLA) foci between MYCN and proteins that are significantly altered in the IMR-32 MYCN interactome following EXOSC10 depletion. Shown are single-cell analyses of one representative replicate for each combination. Lines in boxes depict median and outliers are shown as dots. *P*-values were calculated with Wilcoxon rank-sum test (n=3).

(**C**) Representative images from the quantified PLAs shown in (B). Scale bar: 10 µm.

(**D**) Immunoblot of endogenous MYCN and MAX sedimentation patterns in 10–30% glycerol gradients from IMR-32 shEXOSC10 cells treated with 1 µg/ml Dox for 72 h. Nuclear lysates were treated either with 50 U/ml benzonase or DNase I and RNase inhibitor (50 U/ml each). Note that the lower gradient immunoblot is identical to the one shown in Figure 1D.

**Figure S3: MYCN binds to intronic RNAs independently of DNA binding. Related to Figure 2**.

**(A)** Read distribution of MYCN eCLIPs at the *WDR60*, *SSBP4* and *KCNQ2* genes. IMR-32 cells were treated with 72 h Dox (1 µg/ml) or 1.5 h IAA (100 µM) to knock-down EXOSC10 or EXOSC5, respectively. Control condition treated with EtOH. Input RNA corresponds to the control condition size-matched eCLIP input, U2OS neg. control refers to a MYCN eCLIP performed in U2OS cells that do not express MYCN (n=2 for all eCLIPs).

**(B)** Read distribution of MYCN eCLIPs and ChIP-sequencing (ChIP-Rx) control conditions at the *EIF2B5* and *BIRC5* genes. Same cells and conditions as in (A) (n=2 for all eCLIPs; n=3 for MYCN ChIP-Rx).

**(C)** Number of high-confidence MYCN eCLIP peaks within 1 kb of MYCN ChIP-Rx peak summits. MYCN ChIP-Rx control sample with the highest number of called peaks was used to determine the overlap.

**(D)** (Left) MYCN eCLIP peak distribution at depicted gene regions for indicated conditions. (Right) Distribution of intronic MYCN eCLIP peaks from nearest to farthest from the transcription start site for all shown conditions.

**(E)** Number of UG dinucleotides per MYCN peak in all depicted conditions. Shuffled datasets contain the same number of randomized peaks as the corresponding eCLIP. Lines within box plots indicate median, outliers shown as dots. *P*-values calculated with Wilcoxon rank-sum test.

**Figure S4: Direct binding of MYCN to RNA depends on MYCBoxI. Related to Figure 3**.

(**A**) Coomassie-stained SDS-gel of the recombinant proteins used in RNA-binding assays.

(**B**) Dissociation constant (K_d_) values of His-MBP-tagged MYCN 1-154 construct with depicted series of RNA sequences with variable UG dinucleotide content and indicated length (n=3; n.d.=K_d_ could not be determined).

(**C**) Fluorescence anisotropy (FA) measurement using half-log dilutions from 20 or 10 μM to 0.2 nM of indicated MYCN construct to UG^high^ DNA. Change in anisotropy relative to 5nM FITC-labeled DNA in the absence of protein is plotted relative to each construct. Dots show mean and error bars depict standard deviation (n=3).

(**D**) FA measurement using half-log dilutions of MYCN 372-464 with UG^high^and UG^low^ RNA oligonucleotide. Change in anisotropy relative to 5nM FITC-labeled RNA in the absence of MYCN is plotted relative to MYCN 372-464 concentration. Dots show mean and error bars depict standard deviation (n=3).

**Figure S5: MYCN enhances RNA binding of the nuclear exosome at its binding sites. Related to Figures 5 and 6**.

**(A)** Immunoblot of endogenous MYCN, ZCCHC8 and PABPN1 IPs from IMR-5 cells. GAPDH was used as loading control. The lower MYCN blot image shows a longer exposure of a section of the membrane above containing only the ZCCHC8 and PABPN1 IPs (n=1).

**(B)** Immunoblot of SH-EP MYCNER cells expressing Dox-inducible shRNA targeting SF3B1 (left). Cells were treated for 48 h with 1 µg/ml Dox and 4 h with 100 nM 4-OHT (MYCN ON), where indicated. MYC was used as a control of MYCNER activation, as 4-OHT addition will decrease MYC protein levels. GAPDH was used as loading control (n = 2).

**(C)** Growth curves of SH-EP MYCNER shSF3B1 cells. Cells were treated with 1 µg/ml Dox and 100 nM 4-OHT (MYCN ON) as indicated over the entire measurement time. Dots show mean and error bars depict standard deviation (n=3).

**(D)** Endogenous MYCN and HA-tag immunoprecipitations in IMR-32 cells stably expressing an HA-tagged inactive EXOSC10 mutant (EXOSC10_mut_) (n=2).

**(E)** Trimmed (2%) average density blots of MYCN eCLIP and corresponding size-matched input reads in depicted conditions, centered around EXOSC10_mut_ control (left) and MYCN-depleted (right) high-confidence eCLIP peaks. Only reads in the sense strand are shown. Lines correspond to mean of biological duplicates, shaded regions depict standard error.

**Figure S6: RNA-bound MYCN promotes the degradation of non-polyadenylated intronic RNA. Related to Figure 7**.

(**A**) Trimmed (2%) average density blots of 3’ RNA seq with additional polyadenylation (all RNAs) in IMR-32 shEXOSC10 or shMYCN cells. Reads extend 1 kb upstream and 2 kb downstream of annotated promoter upstream transcript (PROMPT) start sites. Only reads in the sense strand are shown. Reads for each replicate are shown, shaded regions depict standard error. TSS: transcription start site.

(**B**) Lengths of all introns and of introns containing MYCN and EXOSC10_mut_ high-confidence eCLIP peaks in depicted conditions. Median lengths in nt are shown next to each box plot. Lines within box plots indicate median, outliers shown as dots.

(**C**) Log_2_FC comparing EXOSC10 or MYCN depletion to control condition at the 3’ ends of introns bound by MYCN in EXOSC5 AID and shEXOSC10 conditions. Introns are stratified according to length as shown.

