## Supplementary figures and images for "Direct RNA-binding by MYCN mediates feedback from RNA processing to transcription control"

### Figure S1

Figure S1 (Papadopoulos, Ha, Fleischhauer et al.)

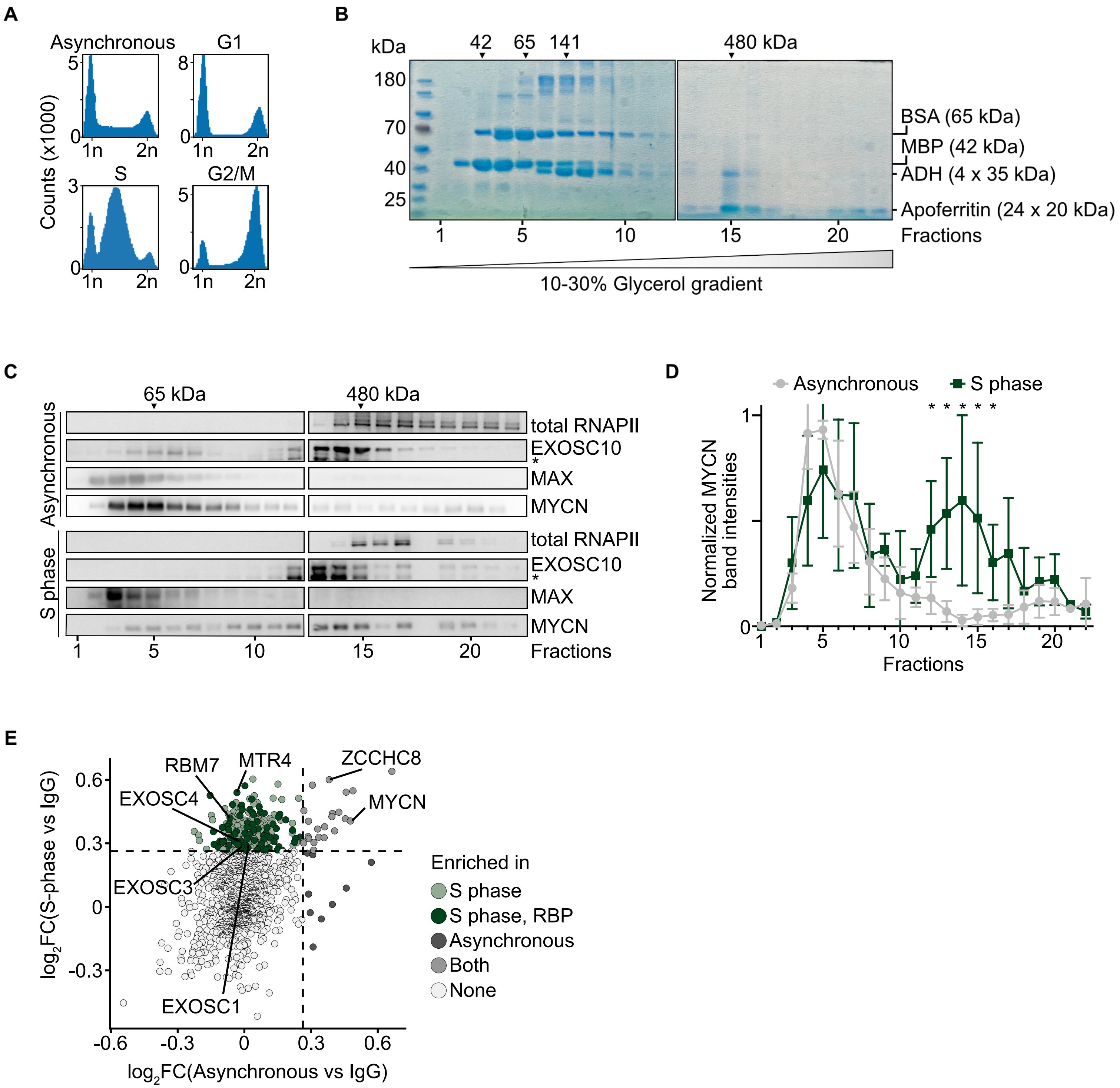

### Figure S2

Figure S2 (Papadopoulos, Ha, Fleischhauer et al.)

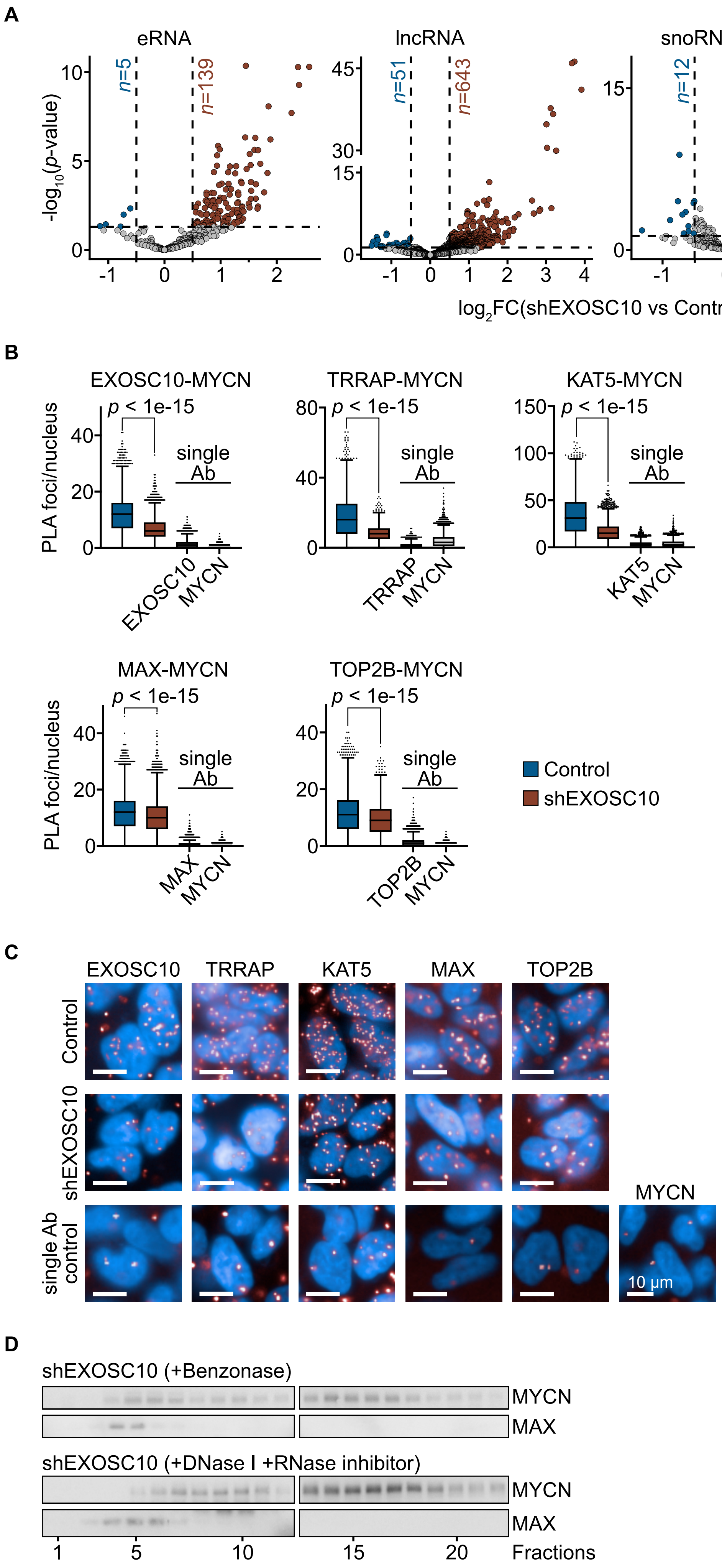

### Figure S3

Figure S3 (Papadopoulos, Ha, Fleischhauer et al.)

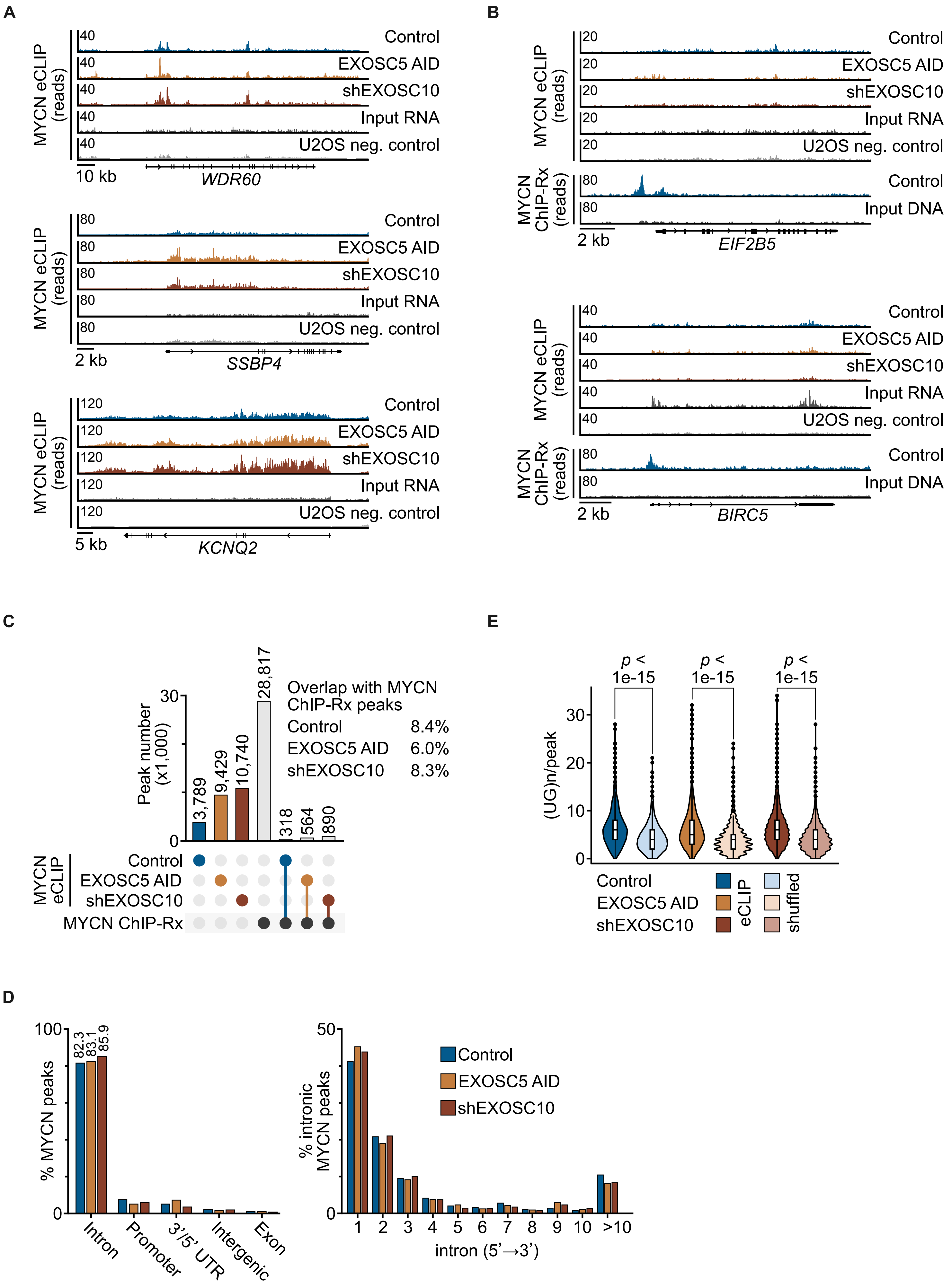

### Figure S4

Figure S4 (Papadopoulos, Ha, Fleischhauer et al.)

A

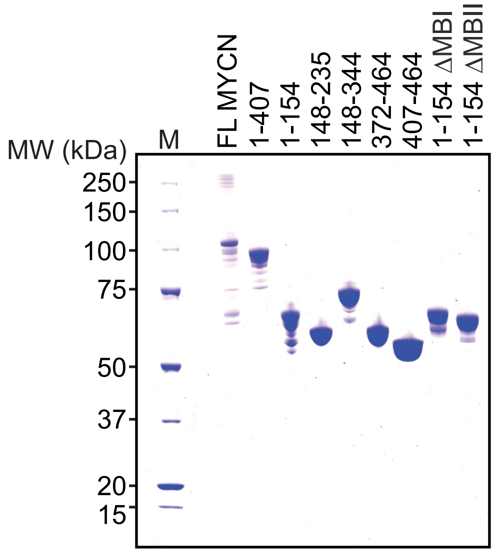


C

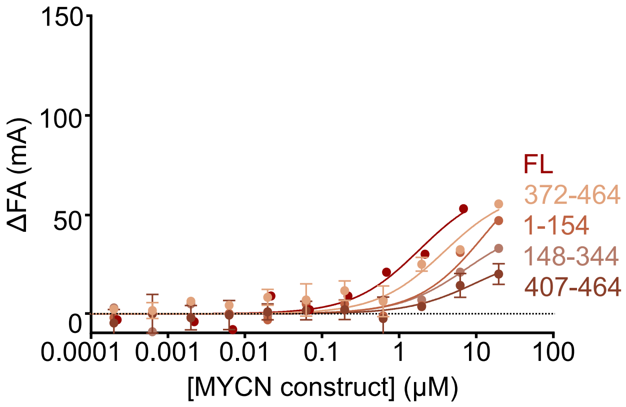


B

| Length | $K_d$ MYCN 1-154 |
|---|---|
| 62 nt | 1.3 (1.1-1.9) |
| 64 nt | n.d. |
| 67 nt | 6.5 (1.6-84.5) |
| 74 nt | n.d. |
| 80 nt | 2.4 (1.2-5.9) |

D

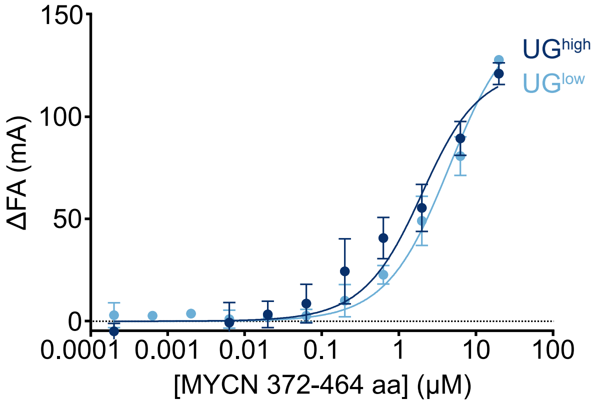

### Figure S5

Figure S5 (Papadopoulos, Ha, Fleischhauer et al.)

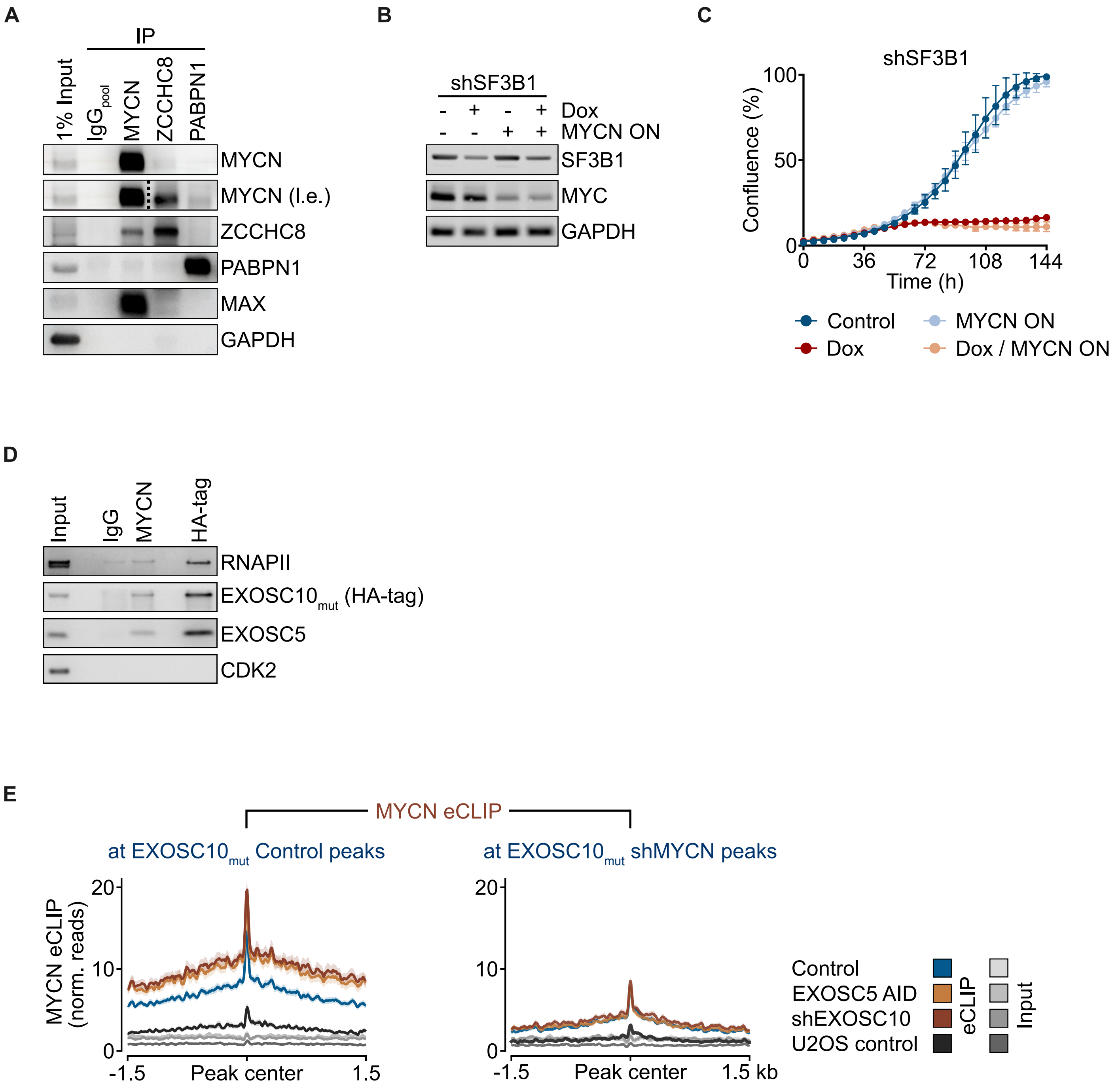

### Figure S6

Figure S6 (Papadopoulos, Ha, Fleischhauer et al.)

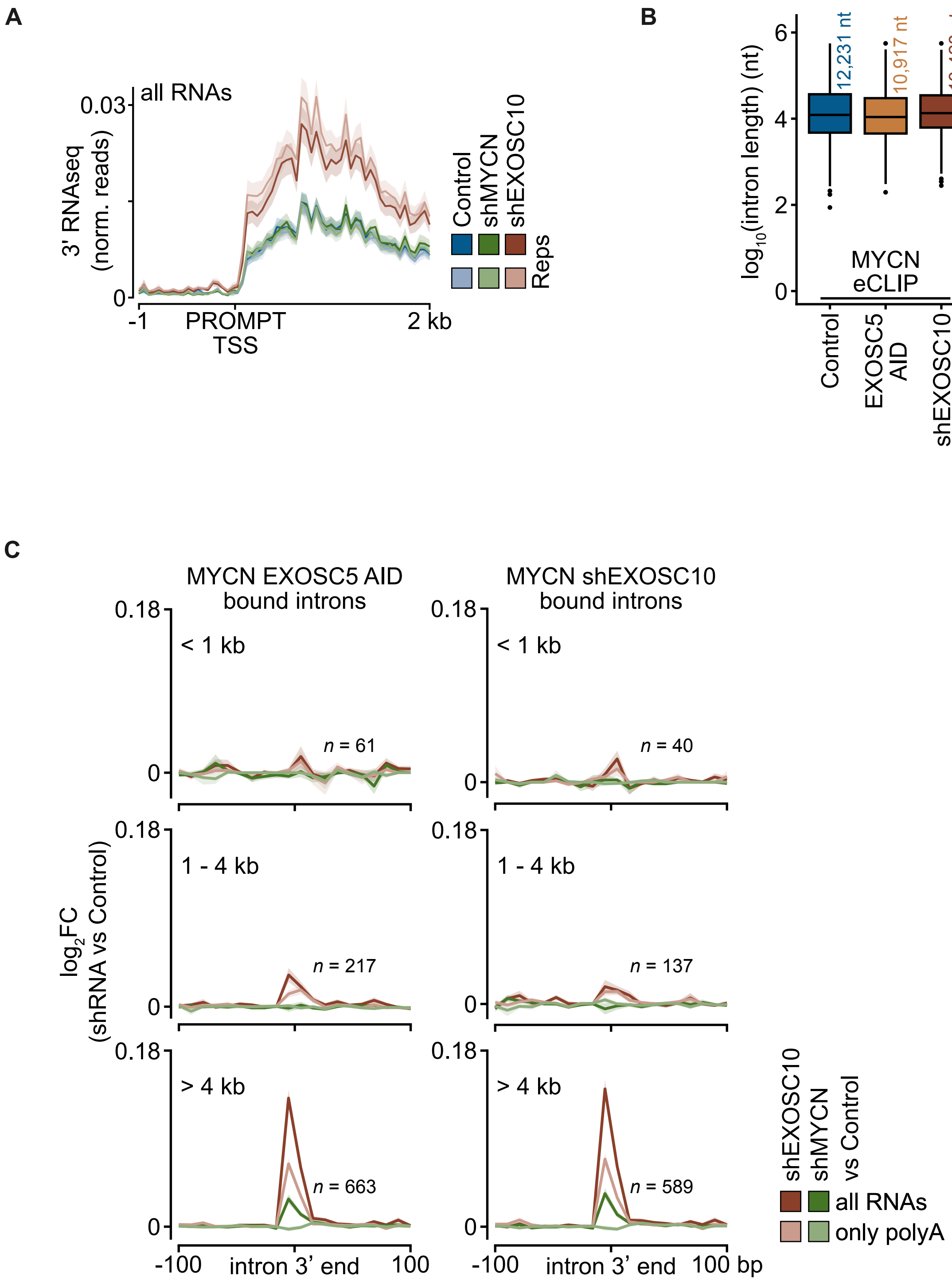
